# *CRY2* missense mutations suppress P53 and enhance cell growth

**DOI:** 10.1101/2021.01.08.425994

**Authors:** Alanna B. Chan, Gian Carlo G. Parico, Jennifer L. Fribourgh, Lara H. Ibrahim, Michael J. Bollong, Carrie L. Partch, Katja A. Lamia

## Abstract

Disruption of circadian rhythms increases the risk of several types of cancer. Mammalian cryptochromes (CRY1 and CRY2) are circadian transcriptional repressors that are related to DNA repair enzymes. While CRYs lack DNA repair activity, they modulate the transcriptional response to DNA damage, and CRY2 can promote SCF^FBXL3^-mediated ubiquitination of c-MYC and other targets. Here, we characterize five mutations in CRY2 observed in human cancers in The Cancer Genome Atlas. We demonstrate that two orthologous mutations of mouse CRY2 (D325H and S510L) accelerate the growth of primary mouse fibroblasts expressing high levels of c-MYC. Neither mutant affects steady state levels of overexpressed c-MYC, and they have divergent impacts on circadian rhythms and on the ability of CRY2 to interact with SCF^FBXL3^. Unexpectedly, stable expression of either CRY2 D325H or of CRY2 S510L robustly suppresses P53 target gene expression, suggesting that this is the primary mechanism by which they influence cell growth.

## Introduction

Circadian rhythms entrain many aspects of physiology to the daily solar cycle (Bass, 2012). Mammalian circadian rhythms are generated by a cell autonomous molecular clock based on a transcription-translation feedback loop (TTFL): a heterodimer of circadian locomotor output cycles kaput (CLOCK) and brain and muscle ARNT-like protein 1 (BMAL1) drives transcription of their own repressors period (PER 1-3) and cryptochrome (CRY1-2). The F-box and leucine-rich repeat proteins 3 (FBXL3) and 21 (FBXL21) are substrate adaptors for S phase kinase-associated protein 1 (SKP1)-cullin 1 (CUL1)-F-box protein (SCF) E3 ubiquitin ligases that stimulate CRY ubiquitination and degradation and contribute to circadian rhythms (Busino et al., 2007; Dardente et al., 2008; Godinho et al., 2007; Hirano et al., 2013; Siepka et al., 2007; Xing et al., 2013; Yoo et al., 2013).

In addition to driving circadian rhythms, clock components transmit temporal information to other physiological processes (Bass, 2012; Hunt and Sassone-Corsi, 2007; Koike et al., 2012; Kowalska et al., 2013). For example, CRY1/2 suppress the activity of several non-circadian transcription factors (Chan et al., 2020; Huber et al., 2016; Jang et al., 2016; Jordan et al., 2017; Koike et al., 2012; Kriebs et al., 2017; Lamia, 2011), thereby influencing their susceptibility to activation in a time of day dependent manner. In the case of c-MYC and Early 2 factor (E2F) family members, this suppression is driven by stimulated proteolysis (Chan et al., 2020; Huber et al., 2016), in which CRY2 acts as a co-factor to recruit c-MYC or E2F family members to the SCF^FBXL3^ complex to promote ubiquitination and subsequent degradation.

Mammalian CRYs evolved from bacterial light-activated DNA repair enzymes also known as photolyases, but lack DNA repair activity and light sensitivity (Partch and Sancar, 2005). Several studies have identified molecular links between CRYs and cancer-related pathways (Chan and Lamia, 2020). The CRY photolyase homology region (PHR) comprises most of the protein, excluding the disordered C-terminal tail (CTT). The PHR includes a flavin adenine dinucleotide (FAD) binding pocket, a secondary pocket (analogous to the antenna chromophore-binding pocket of photolyases (Sancar, 2003)), and a coiled-coil (CC) just upstream of the CTT. FBXL3 and FBXL21 interact with the FAD binding pocket of CRYs; a C-terminal tryptophan in each of these F-box proteins occupies the FAD binding site (Busino et al., 2007; Dardente et al., 2008; Godinho et al., 2007; Hirano et al., 2013; Siepka et al., 2007; Xing et al., 2013; Yoo et al., 2013). Similarly, a tryptophan in the HI loop of CLOCK interacts with the secondary pocket of CRY1 (Michael et al., 2017). Sequence differences between CRY1 and CRY2 surrounding the secondary pocket have a major influence on their differential binding affinity for CLOCK:BMAL1 (Fribourgh et al., 2020; Rosensweig et al., 2018) and their divergent influence on circadian period (Rosensweig et al., 2018). The CC helix of CRY1 interacts with both PER2 and with the TAD of BMAL1 and likely reduces the association of BMAL1 with co-activators (Nangle et al., 2014; Schmalen et al., 2014; Xu et al., 2015). The CTTs in CRY1 and CRY2 are highly divergent both from the C-termini of CRYs in other organisms and from each other. The CTTs contain many phosphorylation sites, which modulate stability (Gao et al., 2013b; Harada et al., 2005; Kurabayashi et al., 2010; Papp et al., 2015), are regulated by DNA damage signaling (Gao et al., 2013b; Papp et al., 2015), and influence circadian timing (Parico et al., 2020).

The circadian and cell cycles influence each other (Bieler et al., 2014; Geyfman et al., 2012; Hong et al., 2014; Kowalska et al., 2013; Matsuo et al., 2003) and several studies have demonstrated that environmental or genetic disruption of circadian rhythms alters cancer development (Pariollaud and Lamia, 2020). We previously demonstrated that genetic deletion of *Cry2* renders cells more susceptible to transformation by diverse oncogenic insults (Huber et al., 2016). Here, we investigate five missense mutations in CRY2 that have each been observed in multiple human tumors, selected based on their predicted impacts on the interactions between CRY2, FBXL3, and c-MYC. Expression of wildtype CRY2, but not of two of the mutant mouse orthologues studied here, suppresses the growth of c-MYC-transformed *Cry2*^−/−^ fibroblasts (Huber et al., 2016). One of the mutants, human CRY2 S532L (mouse CRY2 S510L), exhibits greatly reduced association with FBXL3 as predicted. In contrast, the human CRY2 D347H mutation (mouse CRY2 D325H) does not impact the association of CRY2 with FBXL3 or c-MYC. Instead, we found that it reduces the reduces the ability of CRY2 to interact with and repress the CLOCK:BMAL1 heterodimer, and therefore, it cannot support circadian rhythms in fibroblasts. To gain insight into common effects of the mouse CRY2 D325H and S510L mutations, we globally sequenced RNA prepared from cells expressing wildtype or mutant CRY2 in combination with c-MYC. This unbiased analysis revealed that both CRY2 missense mutations that accelerate transformation by c-MYC robustly reduce the expression of P53 target genes, suggesting that these mutations influence cell growth by suppressing P53.

## Results

### Missense mutation of CRY2 alters cell growth in a context-dependent manner

We used cBioPortal (Cerami et al., 2012; Gao et al., 2013a) (last accessed November 13, 2020) to identify recurrent missense mutations in CRY2 reported in human tumor samples in The Cancer Genome Atlas PanCancer Atlas studies (Figure 1, Figure Supplement 1). We selected five mutations to investigate based on *(i)* their frequency of observation in diverse tumor types (when cBioPortal was first accessed in August 2016), and *(ii)* their surface-exposed locations on the three-dimensional structure of CRY2 in close proximity to the CRY2:FBXL3 interface and the phosphate-binding loop where c-MYC interacts with CRY2 (Figure 1A), suggesting that these mutations may affect the interactions between CRY2, FBXL3, and c-MYC.

**Figure 1.**
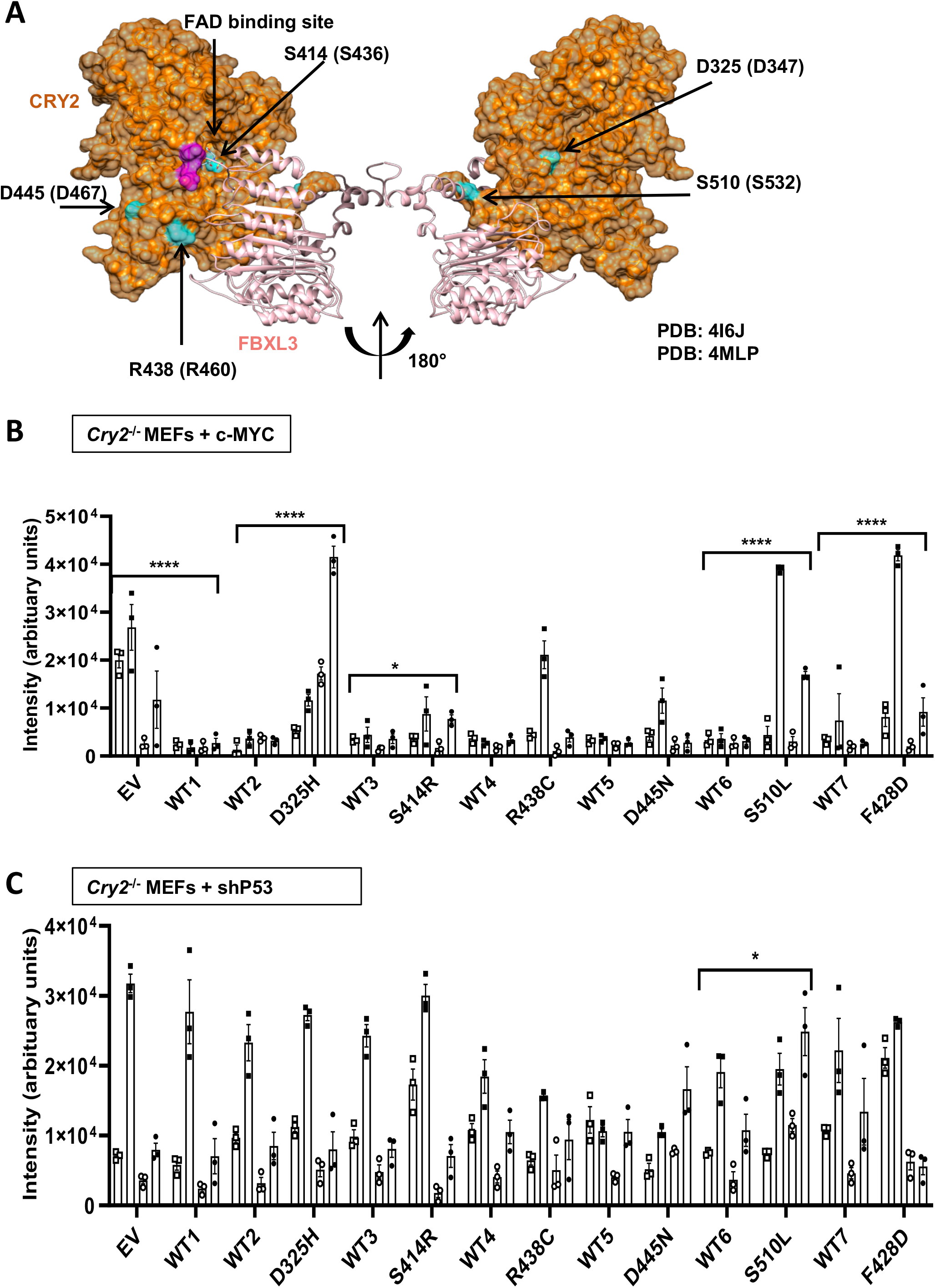
Missense mutations in CRY2 alter cell growth. (A) Three-dimensional structure of mouse CRY2 (orange) and FBXL3 (light pink) (PDB ID: 4I6J). The location and numbering of amino acids orthologous to missense mutations found in The Cancer Genome Atlas (TCGA) are depicted in blue, with the relevant mutations in human CRY2 indicated in parentheses. Magenta shading on CRY2 indicates the phosphate binding loop that interacts with c-MYC. (B,C) Quantitation of crystal violet staining of colonies formed by *Cry2*^−/−^ MEFs stably expressing c-MYC (B) or shRNA targeting P53 (C) after plating at low density. Bars represent mean ± s.e.m. for quantification of staining from four biological replicates each analyzed in triplicate, with individual measurements for each replicate depicted by a unique symbol type: open or filled square, open or filled circle. Each condition was compared to controls that were plated in wells on the same plates (see Figure 1 – Figure Supplement 3 and Figure Supplement 4). *p≤0.05, ****p≤0.0001 by two-way ANOVA with Tukey’s multiple comparison test.

Malignant cells must replicate indefinitely (Hanahan and Weinberg, 2011). To investigate the potential for mutant CRY2 to influence cell proliferation, we monitored cell numbers over ten days after sparsely plating *Cry2*^−/−^ mouse embryonic fibroblasts (MEFs) expressing either c-MYC or shRNA targeting P53 in combination with wildtype or mutant CRY2. Cells expressing elevated c-MYC proliferated more quickly, consistent with the well-established role of MYC in promoting entry into S phase (Kaczmarek et al., 1985). Stable expression of wildtype CRY2 slowed proliferation of *Cry2*^−/−^ fibroblasts expressing c-MYC and tended to slow the growth of P53-depleted cells (Figure 1, Figure Supplement 2). While the impact of CRY2 expression on proliferation was reduced by several of the mutations examined, the high variability and small effect size made it difficult to measure significant differences in this assay. Intriguingly, cells expressing CRY2 S414R seemed to proliferate more slowly in the context of P53 depletion.

Two-dimensional colony formation tests cells’ ability to survive and grow under limited paracrine signaling conditions. We previously demonstrated that *Cry2*^−/−^ fibroblasts form colonies much more efficiently than matched wildtype cells in the context of either *c-MYC* overexpression or depletion of *p53*. Notably, this phenomenon was most robust in primary fibroblasts that were maintained in culture only for very few passages (Huber et al., 2016), perhaps due to accelerated accumulation of DNA damage in *Cry2*^−/−^ cells (Papp et al., 2015). To study the impact of CRY2 missense mutations, we generated *Cry2*^−/−^ fibroblasts stably expressing either c-MYC or shRNA targeting P53 and then further manipulated those lines to stably express wildtype or mutant CRY2. The process of generating the cell lines and growing a sufficient number to plate multiple replicates required performing colony formation assays after 10-12 passages. In this context, we found that wildtype CRY2 consistently suppresses colony growth in c-MYC-expressing *Cry2*^−/−^ MEFs (Figure 1B) but the tendency for CRY2 to suppress colony formation in P53-depleted cells was less robust (Figure 1C). Consistent with our previous studies (Huber et al., 2016), a mutant that abolishes the interaction between CRY2 and FBXL3 (F428D) (Xing et al., 2013) fails to suppress colony growth of c-MYC-expressing cells (Figure 1B). Each of the cancer-associated missense mutants of CRY2 that we examined tended to be somewhat less effective at suppressing colony formation in c-MYC expressing cells compared to wildtype CRY2 but most had a mild impact. Strikingly, mouse CRY2 D325H and S510L consistently failed to suppress colony growth in c-MYC expressing cells (Figure 1B and Figure 1, Figure Supplement 3). In the context of *P53* depletion, expression of CRY2 S510L slightly enhanced colony formation, but neither wildtype nor mutant CRY2 had a major impact on colony formation of P53-depleted cells under the conditions tested (Figure 1C and Figure 1, Figure Supplement 4).

### CRY2 missense mutations alter protein-protein interactions

To determine whether the observed impact of CRY2 missense mutations on cell growth could be explained by altered interaction with SCF complexes and/or with c-MYC, we examined the interactions of wildtype and mutant CRY2 with FBXL3, FBXL21, and c-MYC. Human CRY2 S436R, R460C, and S532L each exhibits decreased interactions with FBXL3 and FBXL21 compared to wildtype CRY2 (Figure 2A,B), though their impact is more subtle than that of the CRY2 F428D mutant, which was established from structural studies to critically disrupt interaction with FBXL3 (Xing et al., 2013) (Figure 2, Figure Supplement 1). None of the mutants consistently affected interactions with c-MYC (Figure 2, Figure Supplement 2).

**Figure 2.**
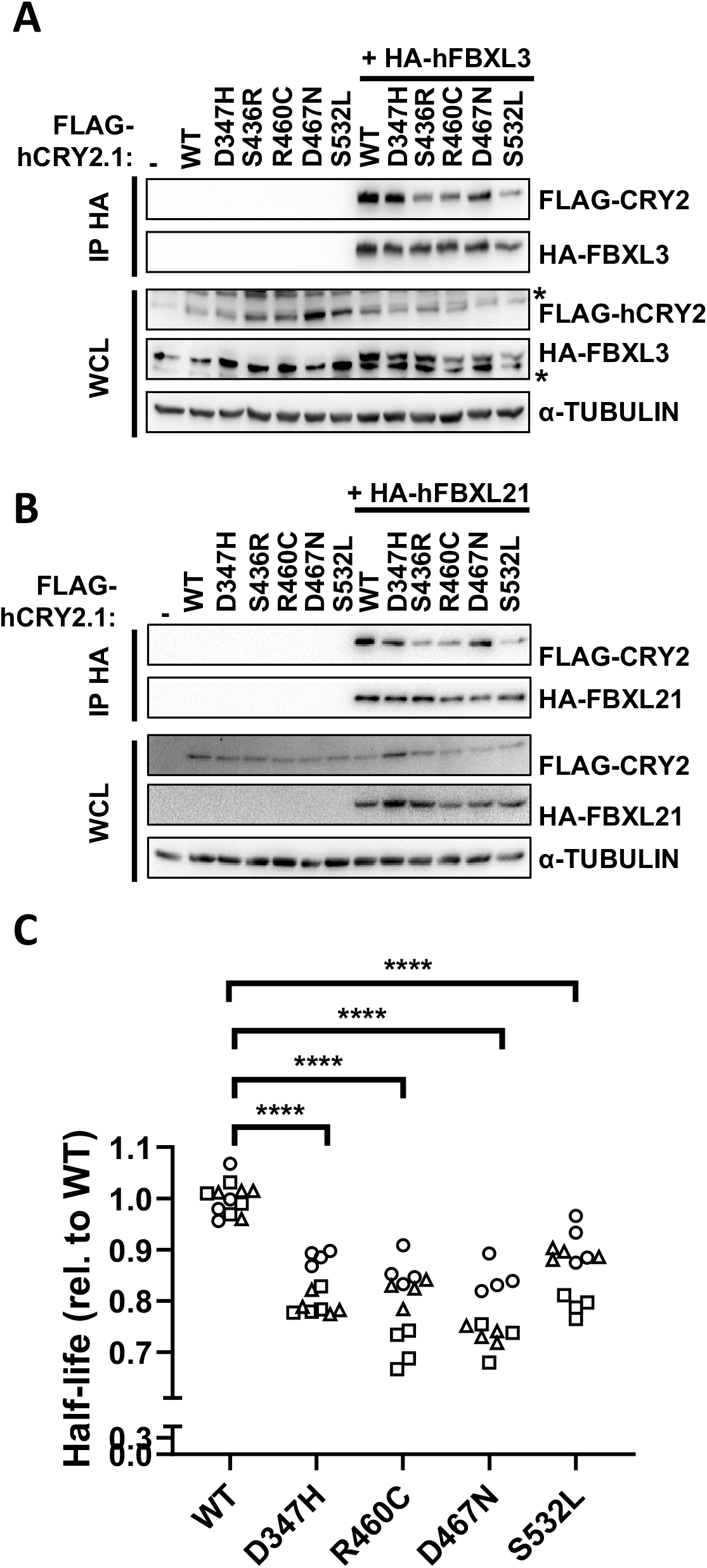
CRY2 S436R, R460C, and S532L exhibit reduced interaction with F-box proteins. (A,B) Proteins detected by immunoblot (IB) following HA immunoprecipitation (IP) or in whole cell lysates (WCL) from HEK293T cells expressing the indicated plasmids with the indicated tags. * denotes non-specific band. (C) Half-life of CRY2-Luciferase fusions proteins containing the indicated missense mutations in CRY2 expressed in AD293 cells. Each symbol represents the half-life calculated from fitting luminescence data recorded continuously after addition of cycloheximide to an exponential decay function. Data represent three independent experimental replicates indicated by different symbols, each performed in quadruplicate. ****p≤0.0001 by two-way ANOVA with Tukey’s multiple comparison test.

Since FBXL3 and FBXL21 are primary determinants of CRY stability (Busino et al., 2007; Dardente et al., 2008; Godinho et al., 2007; Hirano et al., 2013; Siepka et al., 2007; Xing et al., 2013; Yoo et al., 2013), we hypothesized that the half-lives of CRY2 S436R, R460C, and S532L would be increased due to their reduced interaction with FBXL3 and FBXL21. We generated tools to express fusion proteins in which luciferase is appended to the C-terminus of wildtype or mutant CRY2 (CRY2::LUC) in AD293 cells to measure the impact of each mutation on CRY2 stability. Contrary to expectations, each of the missense mutations studied reduces the half-life of the CRY2::LUC fusion protein (Figure 2C and Figure 2, Figure Supplement 3). This assay may not accurately reflect CRY2 half-life under physiological conditions, given that the fusion protein may behave differently than CRY2 alone, as well as the probability that expression of the fusion protein alters the stoichiometry of CRY2, endogenous F-box proteins, and/or other components that regulate CRY2 stability *in vivo*. Nevertheless, we did not observe consistent alterations in the steady state accumulation of either CRY2 or c-MYC under the conditions in which cell growth properties were impacted by mutant CRY2 (not shown).

Within the core circadian clock mechanism, CRYs directly repress CLOCK and BMAL1 in the presence or absence of PERs (Gustafson and Partch, 2015; Ye et al., 2014). Several studies have suggested that disruption of circadian rhythms may contribute to tumor development (Pariollaud and Lamia, 2020). So, we examined whether the five selected tumor-associated CRY2 mutants exhibit altered interaction with core clock components. Indeed, several of the missense mutations under investigation reduced the recruitment of CRY2 to the CLOCK:BMAL1 heterodimer. Most strikingly, the interaction of human CRY2 D347H with CLOCK in the presence of BMAL1 was almost undetectable (Figure 3A). The ability of overexpressed CRY2 to interact with endogenous BMAL1 recapitulated the pattern of interaction with overexpressed CLOCK in the presence of overexpressed BMAL1, suggesting that these mutants may influence the interaction of CRY2 with the CLOCK:BMAL1 heterodimer under physiological conditions (Figure 3, Figure Supplement 1). Importantly, none of the mutations examined had any effect on the interactions of CRY2 with overexpressed PER1 or PER2 (Figure 3, Figure Supplement 2), suggesting that the decreased interaction of CRY2 D347H with CLOCK in the presence of BMAL1 is due to local disruptions in the structure rather than a global alteration of protein fold. The analogous mutations in human CRY1 (D307H) and mouse CRY2 (D325H) similarly impacted their interactions with CLOCK and BMAL1 (Figures 3B and 3C).

**Figure 3.**
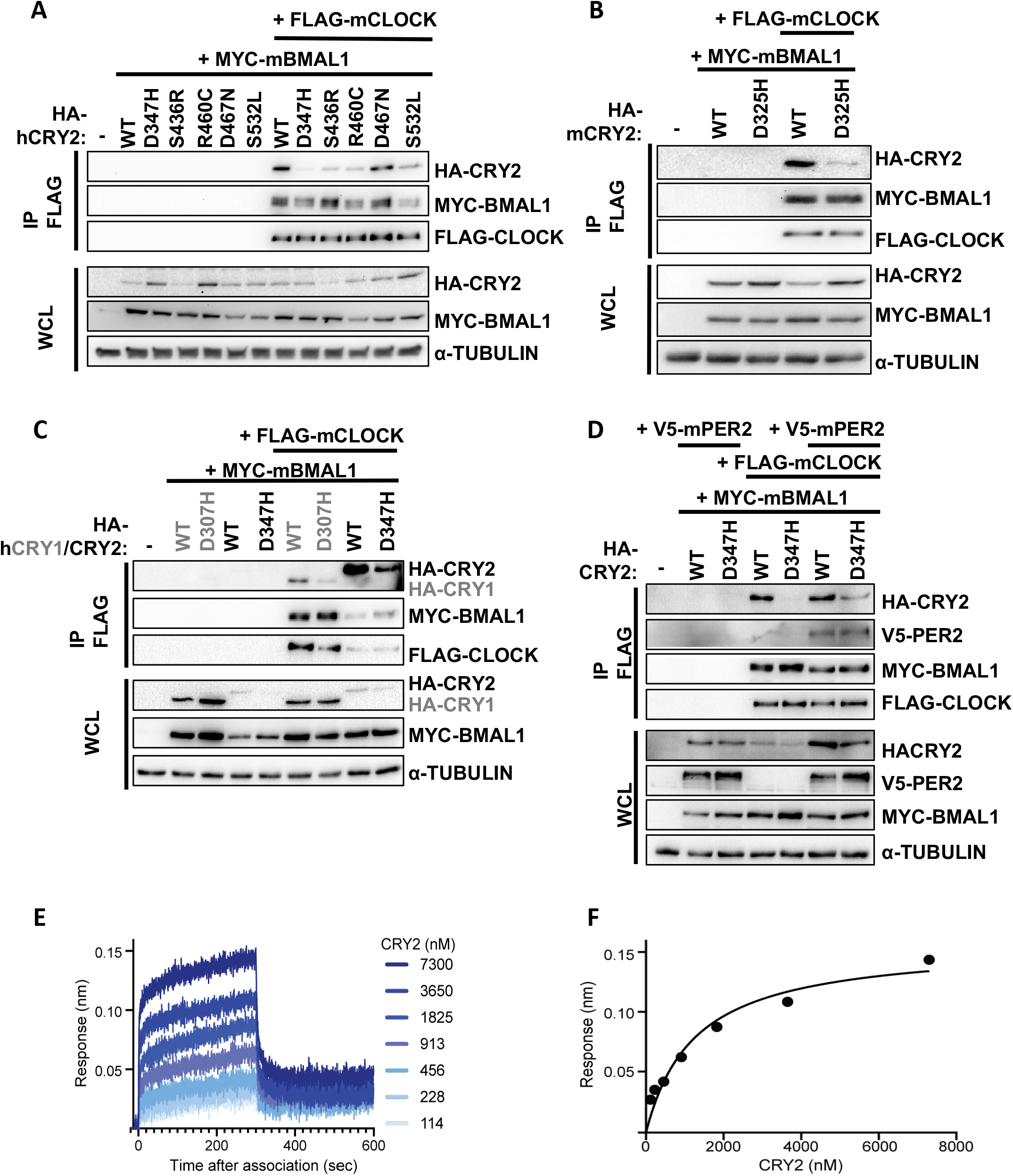
CRY2 D347H, S436R, R460C, and S532L exhibit reduced interaction with CLOCK. (A-D) Proteins detected by immunoblot (IB) following IP of the FLAG tag or in whole cell lysates (WCL) from HEK293T cells expressing the indicated plasmids. (E) Representative BLI binding data of mouse CRY2 D325H PHR domain binding to the CLOCK:BMAL1 PAS domain core (PAS-AB domain heterodimer) immobilized on streptavidin tips. (F) Equilibrium analysis of the BLI data in (E) fit to a one-site binding model. K_D_ = 1.8 ± 0.4μM from two independent BLI experiments.

The interaction of PER2 with CRY2 was previously shown to enhance its ability to co-immunoprecipitate with CLOCK:BMAL1 (Fribourgh et al., 2020; Rosensweig et al., 2018), resulting from an approximately two-fold decrease in the K_D_ of the CRY2 PHR domain for the PAS domain core of CLOCK:BMAL1 *in vitro* (Fribourgh et al., 2020). Consistent with this, co-expression of PER2 enables detection of CRY2 D347H in complex with CLOCK:BMAL1, although the mutant binds less robustly than wildtype CRY2 (Figure 3D). Given that relatively modest changes in affinity impact the ability of CRY2 to co-immunoprecipitate with CLOCK:BMAL1 from cells, we used biolayer interferometry (BLI) to determine how the CRY2 D325H mutation influences the affinity of the mouse CRY2 PHR domain for the PAS domain core of CLOCK:BMAL1 *in vitro* (Figure 3E). The D325H mutation decreased the affinity by 1.5-fold compared to wildtype CRY2 (Fribourgh et al., 2020) for a K_D_ of 1.8 ± 0.4 μM (Figure 3F), helping to explain the reduced interaction of these proteins in cellular extracts.

### CRY2 D347H does not repress CLOCK:BMAL1

Given that CRY2 D347H exhibits reduced interaction with CLOCK in the presence of BMAL1, we predicted that it would be a less effective repressor of CLOCK:BMAL1 transcriptional activity. In U2OS cells, transient overexpression of CLOCK and BMAL1 enhances the expression of luciferase driven by the *Per2* promoter (Figures 4A, 4B). CRY2 dose-dependently decreases the bioluminescent signal, indicating repression of the CLOCK:BMAL1 heterodimer by CRY2 as expected. A previous study demonstrated that CRY2 G351D is unable to repress CLOCK:BMAL1 (McCarthy et al., 2009); we have recapitulated that finding and use CRY2 G351D as a control in our assay. Four CRY2 missense mutants (S436R, R460C, D467N, S532L) examined behave indistinguishably from wildtype CRY2 in this assay. However, consistent with its impact on the interaction of CRY2 with CLOCK and BMAL1 in a co-immunoprecipitation assay, CRY2 D347H does not repress CLOCK:BMAL1 when all three are overexpressed in U2OS cells (Figure 4B).

**Figure 4.**
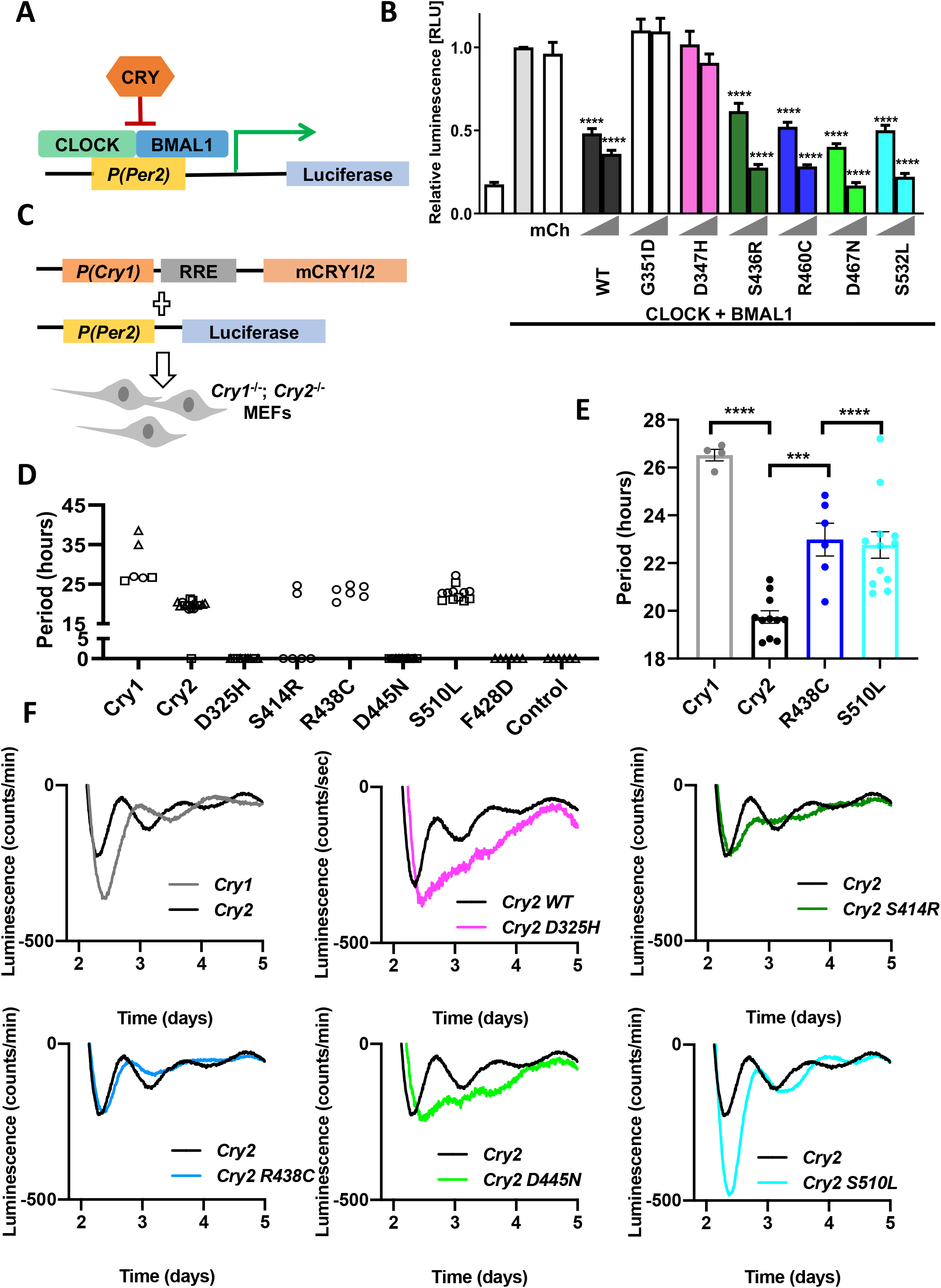
CRY2 D347H neither represses CLOCK/BMAL1 nor supports circadian rhythms. (A) Schematic diagram of luciferase assay performed in (B). (B) Luminescence detected from U2OS cells expressing Per2-Luciferase and the indicated additional plasmids. (mCherry, mCh). Data represented as mean ± SEM, are normalized to gray bar, CLOCK-BMAL1 alone. Triangles on x-axis indicate increasing amounts of CRY wildtype (WT) or mutant transfected, 2 or 5 ng. ****p≤0.0001 by Student’s t-test versus CLOCK-BMAL1 alone in Graph Pad Prism 9. (C) Schematic diagram of rhythmicity rescue assay in Cry1−/−;Cry2−/−MEFs. (D) Periods of Cry1, Cry2 wildtype (WT), or mutant or control (no Cry transfected) across three independent experiments (open circles, triangles, or squares) with six technical replicates in each. Periods calculated using the Lumicycle analysis program using running average background subtraction and fitting to a damped sine wave by least mean squares method (LM sin damped fit). Replicates for which the best fit damped sine had period > 40 hours and/or goodness of fit < 80% were considered arrhythmic and are indicated as 0 on the graph. (E) Periods of the mutants that showed rhythmicity (R438C and S510L). Data represent mean ± SEM of two independent experiments with two-six replicates each as indicated by filled circles in graph. ***p≤0.001, ****p≤0.0001 by one-way ANOVA using Dunnett’s multiple comparisons in Graph Pad Prism 9. (F) Continuous recording of luciferase activity from Cry1−/−;Cry2−/−fibroblasts transfected with Per2-Luciferase and wildtype Cry1 (gray), or wildtype (WT, black) or mutant (as indicated) Cry2. Data represent mean bioluminescence for two-six replicates from a representative of at least two experiments.

To examine whether cancer-associated missense mutations in CRY2 alter the ability of CRY2 to sustain endogenous circadian rhythms, we used the rhythmicity rescue assay developed in (Ukai-Tadenuma et al., 2011). In this assay, arrhythmic *Cry1*^−/−^;*Cry2*^−/−^ MEFs (van der Horst et al., 1999) are transiently transfected with a plasmid encoding destabilized luciferase under the control of the *Per2* promoter, in combination with CRY1 or CRY2 under the control of elements derived from the endogenous *Cry1* promoter and first intron. Expression of wildtype CRY1 (Ukai-Tadenuma et al., 2011) or CRY2 (Rosensweig et al., 2018) under these conditions enables rhythmic expression of the luciferase reporter with long or short periodicity characteristic of *Cry2*^−/−^ or *Cry1*^−/−^ MEFs, respectively. Continuous recording of luciferase activity enables precise measurement of the period of the resulting rhythmic luciferase expression, which allows sensitive detection of perturbations of those rhythms.

Using this system, we found that all of the cancer-associated missense mutations tested impair the ability of CRY2 to sustain short period circadian rhythms of luciferase expression driven by the *Per2* promoter (Figures 4D-F, Figure 4, Figure Supplement 1). Consistent with its inability to repress CLOCK:BMAL1 transcriptional activity, orthologous mouse CRY2 D325H was unable to rescue rhythms. Unexpectedly, CRY2 D445N also failed to support circadian expression of *Per2-luciferase* despite having minimal impact on the interaction of CRY2 with either FBXL3/21 or CLOCK:BMAL1. Expression of CRY2 R438C or S510L resulted in longer periods than those observed with expression of wildtype CRY2. CRY2 S414R sustained detectable rhythms in only one-third of replicates and those rhythms also had a longer period than observed with wildtype CRY2, similar to those supported by CRY2 R438C and S510L.

### CRY2 mutants that fail to suppress colony formation repress P53 target gene expression

Because neither their impact on protein-protein interactions nor their ability to rescue circadian rhythms distinguished the two CRY2 mutants that fail to suppress two-dimensional colony growth in c-MYC expressing fibroblasts (D325H and S510L) (Figure 5, Figure Supplement 1), we took an unbiased approach to examine their influence on global gene expression in those cells. We sequenced RNA from *Cry2*^−/−^ MEFs overexpressing c-MYC and either an empty vector (EV, Control), or wildtype or mutant CRY2 plated at a low or high density. Gene Set Enrichment Analysis (GSEA) (Subramanian et al., 2007) revealed dramatic suppression of a hallmark set of P53 target genes (Liberzon et al., 2015) in cells expressing either CRY2 D325H (FDR = 0.004) or S510L (FDR = 0.01) compared to those expressing wildtype CRY2 (Figures 5A,B,E,F,K,L and Figure 5 Figure Supplement 2). While the effect was less robust than the observed suppression of the P53 pathway, we also measured significantly increased expression of the hallmark hypoxia gene network (FDR = 0.016 or 0.05, Figures 5C,D,G,H,M,N and Figure 5 Figure Supplement 2). Finally, consistent with our demonstration that CRYs cooperate with SCF^FBXL3^ to regulate E2F protein accumulation (Chan et al., 2020), E2F target gene expression was elevated in cells expressing CRY2 S510L (which exhibits decreased interaction with FBXL3), when plated at high density (FDR = 0.21 including only high density samples, Figures 5I,J,O,P,Q and Figure 5 Figure Supplement 2). Taken together, these data suggest that CRY2 D325H and S510L are unable to suppress colony formation of *Cry2*^−/−^ MEFs overexpressing c-MYC due to overlapping functional impairments, including suppression of the P53 transcriptional network.

**Figure 5.**
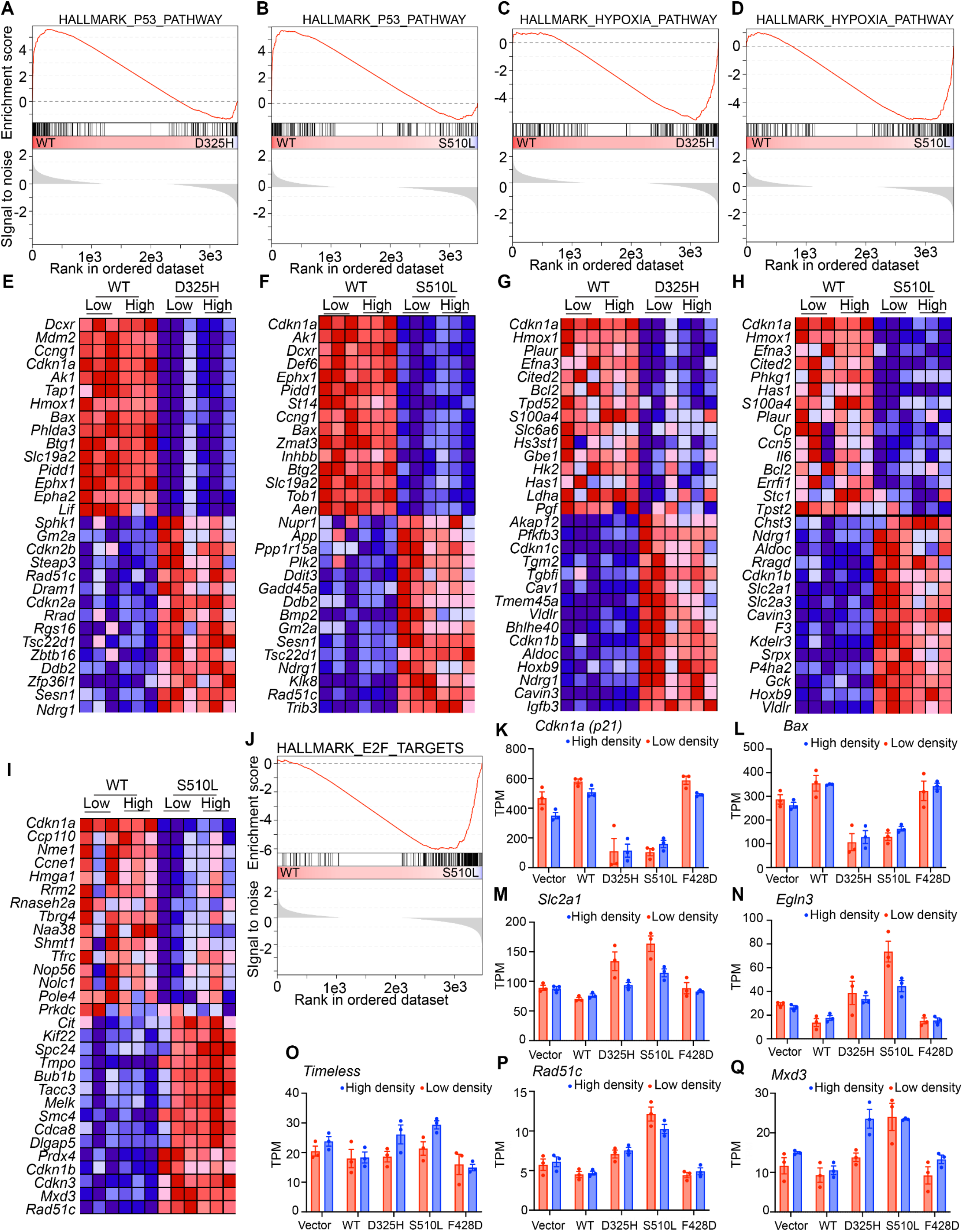
mCRY2 D325H and mCRY2 S510L suppress transcription in the P53 pathway. (A-D, J) GSEA enrichment plots for the hallmark gene sets representing the P53 pathway (A,B), hypoxia pathway (C,D) or E2F target genes (J): The top portion shows the running enrichment score (ES) for the gene set as the analysis walks down the ranked list. The middle portion shows where the members of the gene set appear in the ranked list of genes. The bottom portion shows the value of the ranking metric as you move down the list of ranked genes. The ranking metric measures a gene’s correlation with a phenotype. Here the phenotypes are defined as the cell lines stably expressing wildtype (WT) or mutant (D325H or S510L). (E-I) Heatmaps from RNA-sequencing representing the top 15 up- or down-regulated transcripts of the leading edge that were driving the ES score in (A-D, J). Red, high expression; Blue low expression. (K-Q) Extracted gene expression profiles of the indicated genes from the RNA-sequencing data. Light red, low density plating; light blue, high density plating.

## Discussion

Most missense mutations reported in The Cancer Genome Atlas (TCGA) have unknown functional impact on the cancers in which they are found. Although missense mutations in *CRY2* are rare in TCGA, human genome sequencing data spanning six global and eight sub-continental ancestries (Karczewski et al., 2019) demonstrate that core clock genes, including *CRY2*, are resistant to mutation in healthy somatic tissues, suggesting that recurrent cancer-associated mutations in these genes may be functionally important. Several lines of evidence indicate that *CRY2* may play a role in preventing tumor development. In a variety of cancers, *CRY2* expression is significantly reduced compared to matched normal tissues (Huber et al., 2016; Ye et al., 2018). Furthermore, inactivation or reduced expression of *CRY2* is strongly associated with altered activity of established oncogenic or tumor suppressive pathways (Ye et al., 2018). Here, we demonstrate that two cancer-associated mutations in *CRY2* (D347H and S532L, using numbering for human *CRY2*) prevent CRY2-mediated suppression of colony growth in the context of c-MYC overexpression in *Cry2*^−/−^ fibroblasts.

Several molecular mechanisms have been proposed to explain the apparent inhibition of tumor development by CRY2 (Chan and Lamia, 2020). For example, CRY2 can cooperate with SCF^FBXL3^ to stimulate ubiquitination and subsequent degradation of c-MYC (Huber et al., 2016), the DNA damage activated protein kinase TLK2 (Correia et al., 2019), E2F family members (Chan et al., 2020), and likely other proteins. CRY2 S532L exhibits decreased interaction with FBXL3 and FBXL21, and this likely contributes to the elevated expression of E2F target genes in cells expressing CRY2 S532L. However, disrupted interaction with FBXL3/21 may not be sufficient to prevent CRY2 from suppressing colony formation because the S436R and R460C mutations, which had a similar impact on the ability of CRY2 to interact with FBXL3/21, can suppress colony formation. We cannot exclude the possibility that quantitative differences in the impacts of these mutations on CRY2-FBXL3/21 affinity underlie their diverse functional impacts on cell growth. Conversely, CRY2 D347H fails to suppress colony formation but does not prevent CRY2 from interacting with its F-box protein partners. Thus, disruption of the CRY2-FBXL3 interaction is not required for loss of CRY2 suppression of colony formation in the context of c-MYC overexpression.

CRY2 D347H exhibits reduced interaction with and an inability to repress CLOCK:BMAL1 and cannot sustain circadian rhythms of *Per2-luciferase* expression in *Cry1*^−/−^;*Cry2*^−/−^ fibroblasts. The secondary pocket of CRYs provides an extensive interaction surface for the CLOCK PAS-B domain that plays a critical role in the stable recruitment of cryptochromes to CLOCK:BMAL1 (Fribourgh et al., 2020; Rosensweig et al., 2018). The electrostatic surface of the CRY secondary pocket is mostly negatively charged, while the HI loop of the CLOCK PAS-B domain that docks into this pocket is positively charged (Michael et al., 2017) suggesting that electrostatic attraction could help to support the CRY-CLOCK interaction. Notably, D347 is located on the edge of the secondary pocket of CRY2, providing a rationale for how the mutation from aspartic acid (D) to histidine (H) could contribute to reduced affinity of the CRY2 mutant for CLOCK:BMAL1. Given that differences in the affinity of CRY1 and CRY2 define their ability to stably interact with CLOCK:BMAL1 and repress its activity (Fribourgh et al., 2020; Rosensweig et al., 2018), the D437H mutation represents another example of how a modest change in binding at this critical interface can manifest as a substantial change in CRY function in the clock.

Disrupted circadian rhythms increase cancer risk in humans and increase tumor formation in a variety of genetically engineered mouse models of cancer (Pariollaud and Lamia, 2020). The inability of the mouse ortholog of CRY2 D347H to suppress colony formation in c-MYC expressing fibroblasts supports the contention that circadian rhythms play a protective role against tumorigenesis. However, as for the role of CRY2 in promoting SCF-FBXL3/21 mediated protein turnover, loss of the ability to sustain circadian rhythms is not sufficient to prevent suppression of colony formation since the CRY2 D467N ortholog (D445N) suppresses colony formation but cannot support cell-autonomous rhythms. The inability of CRY2 D445N to sustain circadian rhythms despite its seemingly normal interaction with and repression of CLOCK:BMAL1 suggests that it must disrupt some other aspect of the concerted and periodic interaction with other endogenous proteins (Aryal et al., 2017), appropriate degradation (Godinho et al., 2007; Hirano et al., 2013; Siepka et al., 2007), or other post-translational regulation within the clock mechanism. We cannot exclude the possibility that the impact of the CRY2 mutants examined could differ in the presence of wildtype CRY1 and/or CRY2 both of which are likely present in tumor samples where the mutations occur in a single copy. Taken together, the impacts of the five missense mutations in CRY2 on cell growth, circadian rhythms, and interaction with FBXL3/21 suggest that disruption of core clock function or circadian proteolysis may contribute to enhanced cell growth, but neither is sufficient nor required for loss of growth suppression by CRY2. We measured global RNA expression patterns to identify functional impacts shared by the two mutations with the greatest impact on cell growth. This analysis revealed a striking suppression of a P53 transcriptional network and concomitant elevation of hypoxia target genes in cells expressing mutant CRY2.

We previously demonstrated that CRY1 is stabilized and CRY2 is destabilized in response to DNA damage (Papp et al., 2015). Furthermore, transcriptional activation of at least some P53 target genes is blunted in *Cry2*^−/−^ cells and sustained in *Cry1*^−/−^ cells following DNA damage (Papp et al., 2015). PER2, which cooperates with CRYs to repress transcriptional activation of the CLOCK:BMAL1 heterodimer, stabilizes P53 by inhibiting its MDM2-mediated ubiquitination and degradation (Gotoh et al., 2016; Gotoh et al., 2015; Gotoh et al., 2014). These mechanisms could be at play, but further investigation will be required to understand how CRY2 D325H and CRY2 S510L suppress P53 target gene expression.

Elevated expression of hypoxia-regulated transcripts can enhance cell survival under stressful conditions and contribute to tumorigenesis (Choudhry and Harris, 2018; Semenza, 2013, 2017). We and others recently demonstrated that CRY1 and CRY2 suppress both the protein accumulation and transcriptional activation of HIF1α and HIF2α(Dimova et al., 2019; Vaughan, 2020). HIFs are closely related to CLOCK and BMAL1 (Fribourgh and Partch, 2017) and we and others have shown that BMAL1 dimerizes with HIF1α (Hogenesch et al., 1998; Vaughan, 2020; Wu et al., 2017). Given that the CRY2 D325H mutation reduces its association with and repression of CLOCK:BMAL1, it may also influence repression of HIF1α/BMAL1 by altering protein-protein interactions, although it is unclear whether the interaction of CRYs with HIFs involves the secondary pocket. In contrast, the CRY2 S510L mutation may enhance hypoxia target gene expression by preventing CRY-mediated suppression of HIF1α protein accumulation.

E2Fs are a family of transcription factors that regulate G1 to S phase progression in the cell cycle (Attwooll et al., 2004). The expression of E2F target genes is elevated in cells expressing CRY2 S510L but not in those expressing CRY2 D325H. This distinction likely reflects reduced SCF-FBXL3/21-mediated turnover of E2F family members (Chan et al., 2020) in the presence of CRY2 S510L which exhibits reduced association with FBXL3 and FBXL21. The additional elevation of E2F target genes in combination with suppression of P53 likely contributes to the greater impact of CRY2 S510L in cell growth and colony formation assays.

Over the past decade, several small molecules that target CRYs have been identified and some are under development for therapeutic use in metabolic disease and cancer (Dong et al., 2019; Hirota et al., 2012; Lee et al., 2015; Miller et al., 2020; Oshima et al., 2015). Intriguingly, one of these (TH301) forms a hydrogen bond with the serine in CRY1 analogous to CRY2 S414 and selectively increases the half-life of CRY2 (and not of CRY1) presumably by reducing its interaction with FBXL3 (Miller et al., 2020). The disordered C-terminal tail (CTT) of CRY2 was required for the impact of TH301 on CRY2 stability suggesting that an intramolecular interaction between the CTT and the globular photolyase homology region (PHR) which contains the FAD binding pocket where S414 is located. The apparently greater impact of the CRY2 mutant (D325H) to decrease interaction with CLOCK:BMAL1 in cells compared to the modest difference in affinity between the wildtype and D325H mutant CRY2 PHR for the BMAL1:CLOCK PAS domain core *in vitro* may also be explained by participation of the CTT. Improved understanding of the molecular mechanisms connecting circadian clocks, and CRYs in particular, to cancer-driving networks will inform efforts for further development of CRY-modulating compounds and other strategies to combat the increased incidence of metabolic disease, cancer, and other pathologies associated with chronic circadian disruption.

## Materials and Methods

### Cell culture

HEK 293T and U2OS cells were purchased from the American Type Culture Collection (ATCC). *Cry1*^−/−^; *Cry2*^−/−^ mouse embryonic fibroblasts (MEFs) were generated by Andrew Liu and Hiroki Ueda (Ukai-Tadenuma et al., 2011) and were gifted to us. AD293 cells were gifted to us by Dr. Steve Kay. Cell culture methods were the same as in (Chan et al., 2020). FBS (Thermo Fisher cat # 16000044). All transfections were carried out using linear polyethylenimine (PEI; Polysciences cat. #23966-2) by standard protocols.

### Immunoprecipitation and western blotting

Methods were the same as in (Chan et al., 2020). Antibodies for Western Blots were anti-FLAG polyclonal (Sigma, St. Louis, MO, cat # F7425), anti-V5 polyclonal (Bethyl Labs, Montgomery, TX, cat # A190–120A), anti-HA polyclonal (Sigma, St. Louis, MO, cat # H6908), anti-α-TUBULIN (Sigma, St. Louis, MO, cat # T5168), anti-LAMIN A (Sigma, St. Louis, MO, cat # L1293), CRY1-CT and CRY2-CT as described in (Lamia et al., 2011), anti-c-MYC (Abcam cat # ab32072), anti-P53 (Leica biosystems, Wetzlar, Germany, cat # CM5), anti-MYC (for MYC tag) (Sigma, St. Louis, MO, cat # C3956).

### Plasmids

pcDNA3-2xFlag-mCRY2, pcDNA3-HA-FBXL3, pcDNA3-human c-MYC-V5 are as described previously (Huber et al., 2016). FBXL3 was replaced with human FBXL3 cDNA and human CRY2 cDNA was cloned in to a pCDNA3.1-based HA or 2xFLAG-epitope tagged vector using standard methods. CRY2/CRY1 TCGA mutations were made using Q5 Site-Directed Mutagenesis (New England Biolabs Inc., Ipswich, MA, cat # E0554S). psPAX plasmid (Addgene, Watertown, MA, plasmid 12260) and pMD2.G plasmid (Addgene. Watertown, MA, plasmid 12259) deposited by Dr. Didier Trono, and used for infection, were purchased from Addgene. Watertown, MA. pCL-AMPHO was purchased from Novus Biologicals Littleton, CO (cat # NBP2-29541). MLH-shLuciferase, MLH-shP53 pBABE-Puro, pWZL hygro HRAS^V12^ were gifts from Dr. Tyler Jacks. pcDNA3-2xFlag-mBMAL1, pcDNA3-2xFlag-CLOCK, and pGL2-Per2-luciferase were gifts from Charles Weitz, Department of Neurobiology, Harvard Medical School, Boston. pCMX β-galactosidase was a gift from Dr. Ron Evans, Gene Expression Laboratory, The Salk Institute for Biological Studies, La Jolla. pMU2-P(Cry1)-(intron336)-Cry1/2-Myc, modified with a C-terminal MYC tag was gifted to us by Dr. Carla Green as in (Rosensweig et al., 2018) and the modified using Q5 Site-Directed Mutagenesis (New England Biolabs Inc., Ipswich, MA, cat # E0554S) to create the CRY2 TCGA mutants. p3XFLAG-CMV-14-CRY1-LUC was gifted to us from Dr. Tsuyoshi Hirota as in (Hirota et al., 2012) and modified using Q5 Site-Directed Mutagenesis (New England Biolabs Inc., Ipswich, MA, cat # E0554S) to replace CRY1 with human CRY2 and the CRY2 TCGA mutants.

### Generation of viruses and stable cell lines

Lentiviral or retroviral shRNA or stable overexpression constructs were produced by transient transfections in HEK293T cells using psPAX, PMD2.G, or pCL-AMPHO packaging plasmids for virus generation. Viral supernatants were harvested 48hours after transfection and then filtered through a 0.45μm filter. Viral supernatant was combined with media and 6μg/mL polybrene (Sigma, St. Louis, MO, cat # 107689-10G) and then added to parental cell lines. After 16hrs, the media was changed into selection media – containing 2μg/mL puromycin (Sigma, St. Louis, MO, cat # P9620-10ML), 75μg/mL hygromycin (Sigma, St. Louis, MO, cat # H3274-1G), 5μg/mL blasticidin (Invivogen, San Diego, CA, cat # ant-bl-1). Selection media was replaced every 2-3 days until selection was complete (as judged by death of mock-infected cells, 4 days to 2 weeks).

### Proliferation and 2-D colony formation assays

For proliferation assays, cells were plated at 5000 cells/well in triplicate in a 6-well plate and counted every 2 days for 10 days via hemocytometer. Two independent experiments were performed. For 2-D colony formation assays, cells were plated 150 cells/well of a 6-well plate in triplicate and grown for 11-18 days prior to fixing and staining with crystal violet. To stain, media was aspirated, washed with 1xPBS, then fixed with 100% methanol (Sigma, St. Louis, MO, cat # 179337-4×4L) for 10min and then removed, then stained with 20% methanol and 0.05% crystal violet (Sigma, St. Louis, MO, cat # C3886-25 g) for 20min and then removed, plates were rinsed in DI-H2O and imaged. Three independent experiments were performed.

### Real-time bioluminescence rhythmicity rescue assays

Assays were performed as in (Rosensweig et al., 2018) with some modifications: Cells were transfected with 6ug Per2-luciferase and 150ng of the respective pMU2-P(Cry1)-(intron336)-Cry1/2-Myc or pMU2-P(Cry1)-(intron336)-Cry2-withTCGA mutations-Myc plasmids. Data was analyzed using the Lumicycle Analysis software package. The first two days of recording were discarded, and period, phase, amplitude, and damping rate were calculated using LM fit damped sin wave based on running average fit for each plate of cells. As in (Rosensweig et al., 2018), rescues were considered arrhythmic of the goodness of fit for the damped sin wave was <80% additionally we considered rescues arrhythmic if they had periods of greater than 40 hours. Baseline subtracted data was corrected for background noise as in (Rosensweig et al., 2018).

### Protein degradation assay

Assay was performed as in (Hirota et al., 2012) using AD293 cells with slight modifications. AD293 cells were transiently transfected using PEI. 24hrs after transfection, the media was changed to selection media 750μg/mL Neomycin (G418 Sulfate) (Mirus Bio, Madison, WI, cat # MIR5920). Media was changed ever 2-3 days until selection was complete (as judged by death of mock-infected cells, 4 days to 2 weeks). The stable cell lines were plated at 2×10^6^cells/ 35mm cell culture dish (VWR, Radnor, PA, cat # 82050-538). After 24hrs, media was changed into luciferase recording media (DMEM, 5% FBS, 1% penicillin-streptomycin, 15 mM Hepes, pH 7.6, and 100 μM D-luciferin) for 1 hour then cycloheximide (CHX) (Fisher Scientific, Hampton, NH, cat # 50255724) was added at a final concentration of 100μg/ml diluted in serum free media.

Plates were sealed with vacuum grease (Dow Corning high vacuum grease; VWR, Radnor, PA, cat #59344-055) and glass cover slips (Fisher Scientific Hampton, NH, Cat # 22038999; 40CIR-1,) and placed into the Lumicycle 32 from Actimetrics, Inc. (Wilmette, IL). Data were recorded for 24hrs using the Actimetrics Lumicycle. Data Collection software and analyzed/exported using Actimetrics Lumicycle Analysis program. Half-life was obtained by one phase exponential decay fitting with Prism software (GraphPad Software, Graphpad Holdings, LLC, San Diego, CA).

### Luciferase Assays

Luciferase assays were performed with U2OS cells as in (Kriebs et al., 2017). U2OS cells were plated at 10,000 cells/well in 96-well plates with six replicates per condition After 8 hr, U2OS cells were transfected with 35 ng pGL2-Per2-luciferase, 5 ng β-galactosidase, 0.5 ng BMAL1, 1.5 ng CLOCK, and 0.01–0.05 ng CRY2/mutants/mCherry. All low-concentration plasmid dilutions were prepared fresh immediately before transfection. The following day, medium was replaced. The following day luciferase activity was measured using the britelite plus Reporter Gene Assay System (Perkin-Elmer, Waltham, MA, cat. #6066761) according to the manual, followed by measuring β-galactosidase activity measurement using 2-Nitrophenyl β-D-galactopyranoside (Sigma, St. Louis, MO, cat #N1127) as substrate.

### Mice

*Cry1*^−/−^; *Cry2*^−/−^ mice from which *Cry2*^−/−^ primary MEFs were derived were from Dr. Aziz Sancar (Thresher et al., 1998). All animal care and treatments were in accordance with Scripps Research guidelines and regulations for the care and use of animals. All procedures involving experimental animals were approved by the Scripps Research Institutional Animal Care and Use Committee (IACUC) under protocol #10-0019.

### RNA-sequencing

*Cry2*^−/−^ primary MEFs stably overexpressing c-MYC and EV, WT, D325H, S510L, or F428D were plated in biological triplicate, two wells per replicate, in 6-well plates at two plating densities – 150 cells/ well (low) or 500 cells/ well (high). The cells from two wells were combined and collected for RNA extraction 11 days (high) and 14 days (low) after plating. RNA was extracted using RNeasy mini kit (Qiagen, Hilden, Germany, cat # 74106) according to the manufacturer’s protocol. The QIAGEN RNase-Free DNase Set was used for on-column DNA digestion according to the manufacturer’s protocol (Qiagen, Hilden, Germany, cat # 79256). Total RNA samples were sent to BGI Group, Beijing, China, for library preparation and sequencing. Reads (single-end 50bp at a sequencing depth of 20 million reads per sample) were generated by BGISEQ-500.

### RNA-seq analysis

Kallisto (https://pachterlab.github.io/kallisto/) was used to align to the reference transcriptome (ftp://ftp.ensembl.org/pub/current_fasta/mus_musculus/cdna/) and estimate transcript abundance. GenePattern (https://www.genepattern.org/use-genepattern), Gene Set Enrichment Analysis (GSEA) (Subramanian et al., 2007), and differential expression analysis (DESeq2) (using R (https://www.r-project.org/)) were used to analyze the RNA-seq data for differentially expressed genes and to find suppressed or enriched gene sets across samples.

### Protein expression and purification

His-tagged mouse CRY2 D325H PHR domain (residues 1-512) was expressed in Sf9 suspension insect cells (Expression Systems, Davis, CA) infected with a P3 baculovirus at 1.2 × 10^6^ cells per milliliter and grown for 72 hours at 27°C with gentle shaking. Cells were centrifuged at 4°C for 4000 × rpm for 15 minutes and resuspended in 50 mM Tris buffer pH 7.5, 300 mM NaCl, 20 mM imidazole, 10% (vol/vol) glycerol, 0.1% (vol/vol) Triton X-100, 5 mM β-mercaptoethanol and EDTA-free protease inhibitors (Pierce, Waltham, Massachusetts,). Cells were lysed using a microfluidizer followed by sonication with a ¼” probe on ice for 15 seconds on, 45 seconds off for three pulses at 40% amplitude and lysate was clarified at 4°C for 19,000 rpm for 45 minutes. CRY2 protein was isolated by Ni^2+^-nitrilotriacetic acid (Ni-NTA) affinity chromatography (Qiagen, Hilden, Germany) following elution with 50 mM Tris buffer pH 7.5, 300 mM NaCl, 250 mM imidazole, 10% (vol/vol) glycerol, and 5 mM β-mercaptoethanol. The protein was further purified by HiTrap HP cation exchange chromatography (GE Healthcare, Chicago, IL) and Superdex 75 gel filtration chromatography (GE Healthcare, Chicago, IL) into 20 mM HEPES buffer pH 7.5, 125 mM NaCl, 5% (vol/vol) glycerol, and 2 mM tris(2-carboxyethyl)phosphine (TCEP). The CLOCK:BMAL1 PAS domain core was purified and biotinylated as reported in (Fribourgh et al., 2020).

### Bio-layer interferometry

BLI experiments were performed using an 8-channel Octect-RED96e (ForteBio, Fremont, CA) with a BLI assay buffer of 20 mM HEPES buffer pH 7.5, 125 mM NaCl, 5% (vol/vol) glycerol and 2 mM TCEP. All experiments began with a reference measurement to establish a baseline in BLI assay buffer for 120 seconds. Next, 1.5 μg/mL biotinylated CLOCK:BMAL1 PAS-AB was loaded on streptavidin tips for 300 seconds at room temperature. Subsequently, a 360 second blocking step was performed with 0.5 mg/mL BSA, 0.02% (vol/vol) Tween, 20 mM HEPES buffer pH 7.5, 125 mM NaCl, 5% (vol/vol) glycerol and 2 mM TCEP. Association was measured for 300 seconds with 7 concentrations in a serial dilution of the CRY2 mutant in blocking buffer, and then dissociation was measured for 300 seconds in blocking buffer. Each experiment was repeated with tips that were not loaded with CLOCK:BMAL PAS-AB to provide a reference for non-specific binding to the tip and two independent replicates of the BLI experiment were done. Before fitting, all datasets were reference-subtracted, aligned on the y-axis and for interstep correction through their respective dissociation steps according to the manufacturer’s instructions (ForteBio, Fremont, CA). Data were processed in steady state mode to extract concentration-dependent changes of CRY2 binding to CLOCK:BMAL1 and fit to a one-site binding model using Prism 8.0 (Graph Pad, San Diego, CA).

## Contributions

A.B.C. designed and performed experiments, analyzed data, generated figures, performed GSEA and DESeq2 RNA-sequencing analysis, and wrote the original draft of this paper. BGI performed library preparation and RNA-sequencing. M.J.B aligned the RNA-sequencing data to the mouse transcriptome, performed GSEA, and generated heatmaps. L.H.I. provided R code to run DESeq2 analysis. G.C.G.P and J.L.F. purified recombinant proteins and collected *in vitro* binding data with supervision from C.L.P. K.A.L. performed GSEA, conceived and supervised the study. A.B.C and K.A.L wrote and edited the paper. All authors edited and approved of the manuscript.

## Acknowledgements

This work was supported by NIH grant CA211187 and by a gift from the Brown Foundation for Cancer Research to K.A.L. and NIH grant GM107069 to C.L.P. G.C.G.P. was supported by an HHMI Gilliam fellowship with support from the UCSC Graduate Division. J.L.F. was supported by a UCOP and UCSC Chancellor’s Postdoctoral Fellowship. We thank Drs. Hiroki Ueda, Carla Green, Tsuyoshi Hirota, Clark Rosensweig, Anne-Laure Huber, Megan Vaughan, Anna Kriebs, Marie Pariollaud, and Ms. Rebecca Mello for helpful discussions and/or gifting us critical reagents. The results published here are in whole or part based upon data generated by the TCGA Research Network. We thank Toni Thomas, Judy Valecko, and Yolanda Slivers for administrative assistance.

---

**Figure 1 - Figure Supplement 1.**
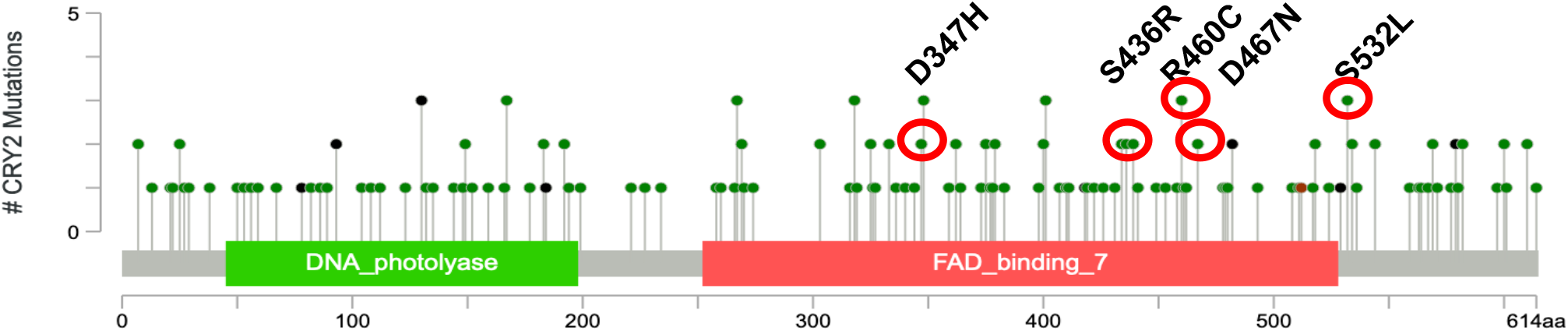
Lollipop diagram from cbioPortal (last accessed November 13, 2020) indicating location and frequency of mutations in CRY2 in tumor samples included in The Cancer Genome Atlas (TCGA). The missense mutations we chose to study in this paper are circled in red.

**Figure 1 - Figure Supplement 2.**
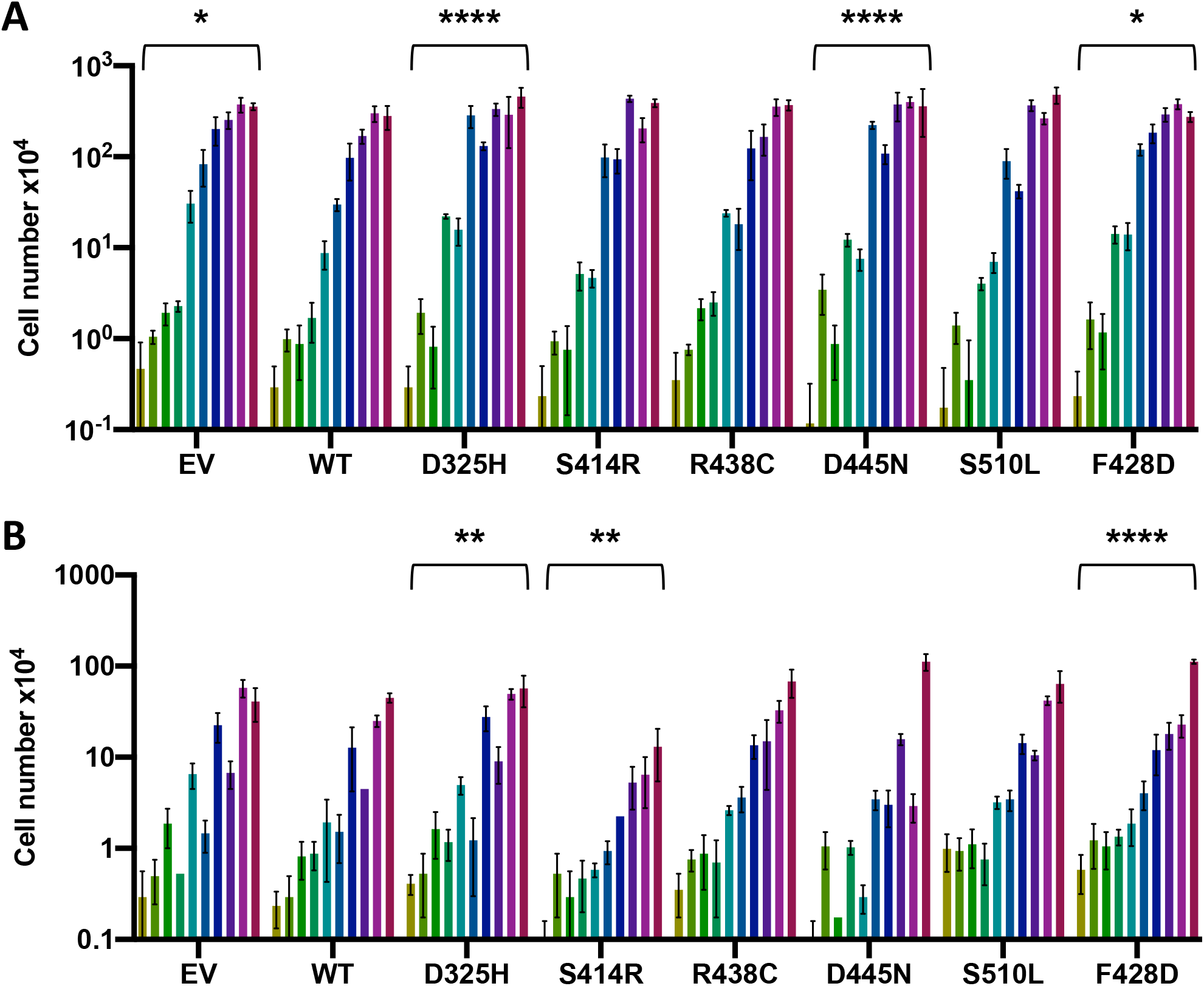
(A,B) Proliferation data expressed as cell number x104 across ten days starting at day two for *Cry2^-/-^*MEFs stably expressing c-MYC (A) or shRNA targeting P53 (B) with additional stable expression of empty vector (EV, control), or wildtype or mutant CRY2 as indicated. Data represent mean ± SEM for two independent experiments (adjacent bars) with three technical triplicates each. *p≤0.05, **p≤0.01, ****p≤0.0001 by two-way ANOVA with Tukey’s multiple comparison test.

**Figure 1 - Figure Supplement 3.**
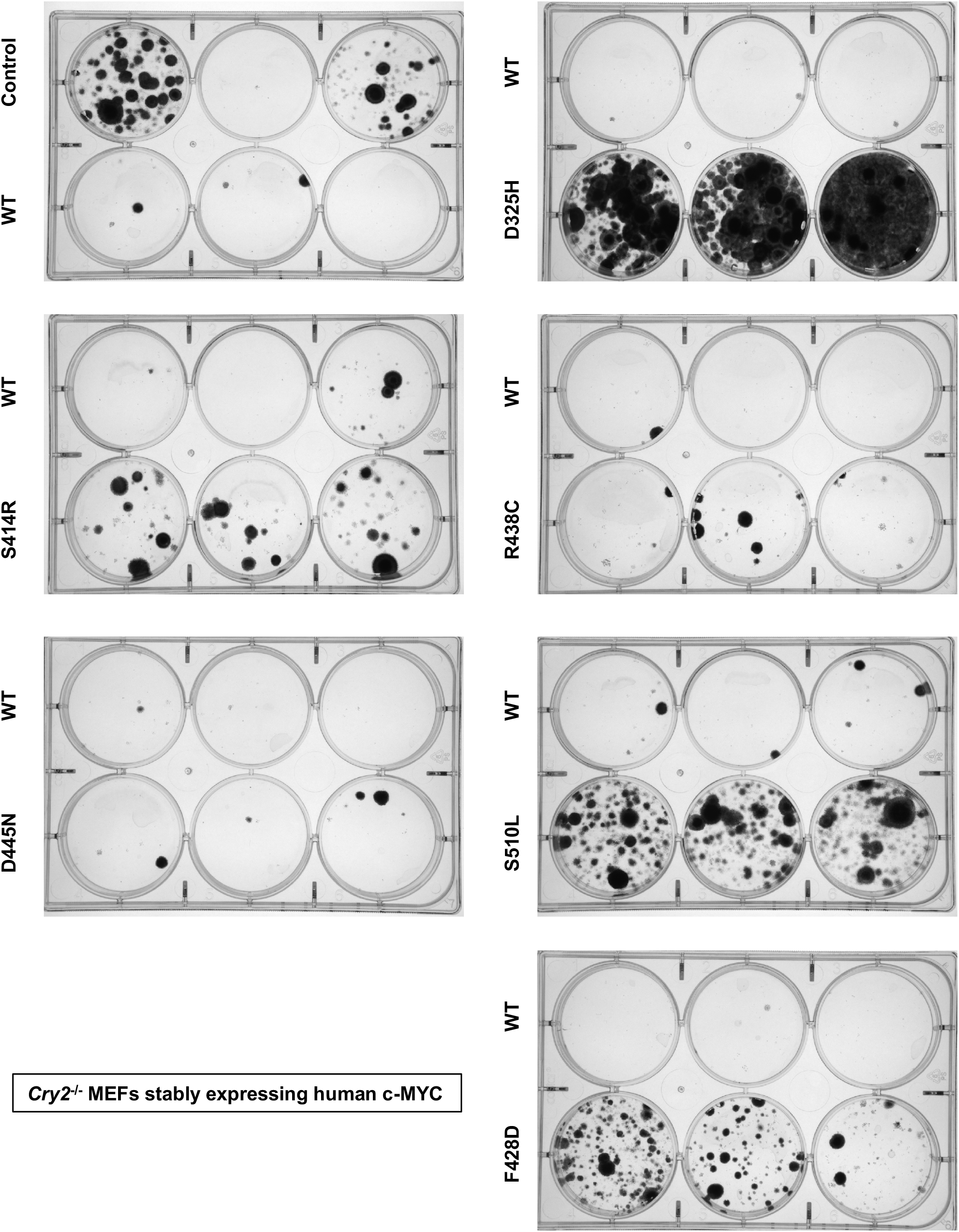
Raw images of one of four biological replicates in Figure 1B.

**Figure 1 - Figure Supplement 4.**
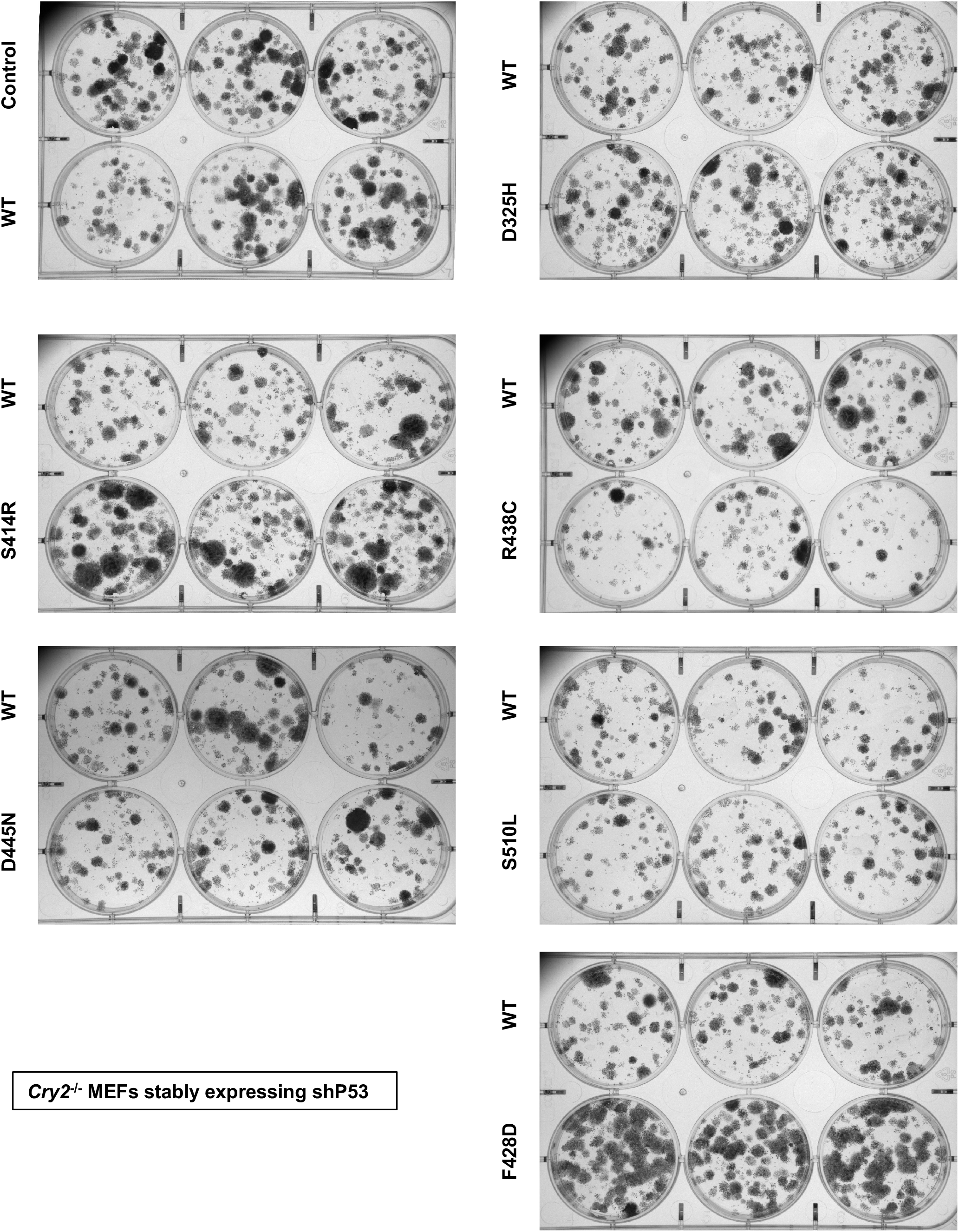
Raw images of one of four biological replicates in Figure 1C.

**Figure 2 - Figure Supplement 1.**
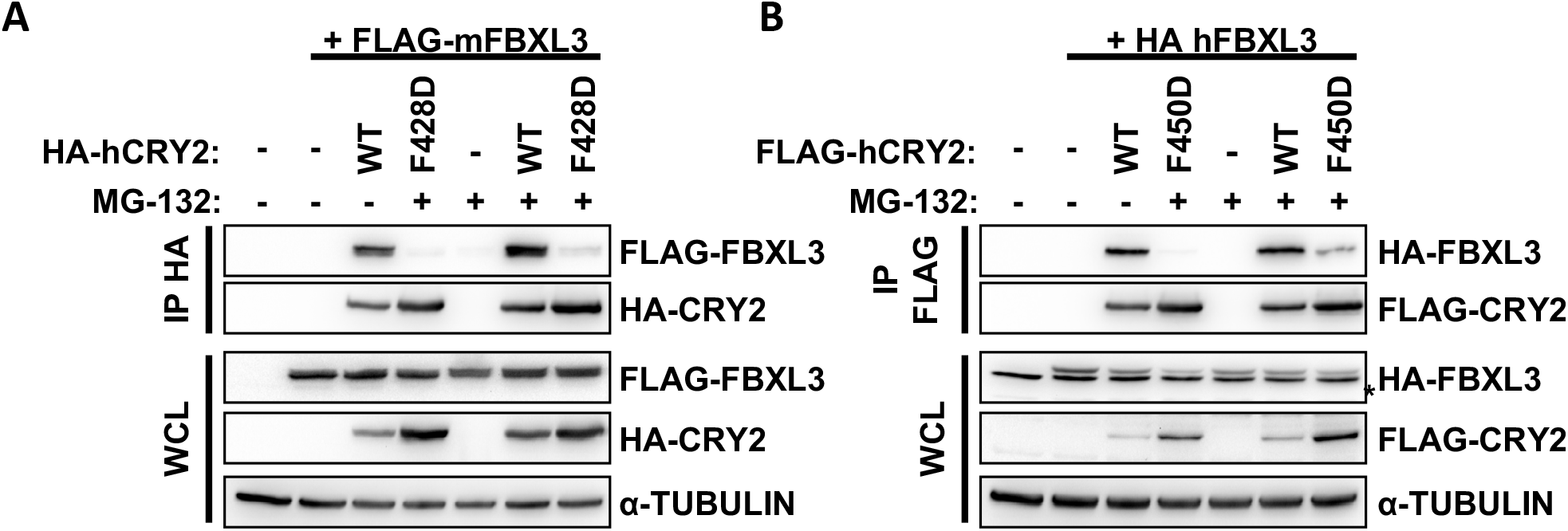
(A,B) Proteins detected by immunoblot (IB) following immunoprecipitation (IP) of HA (A) or FLAG (B) epitope tags from whole cell lysates (WCL) of HEK293T cells expressing the indicated plasmids. Samples were treated with vehicle (-, DMSO) or 10#x03BC;M MG-132 (+) for four hours.

**Figure 2 - Figure Supplement 2.**
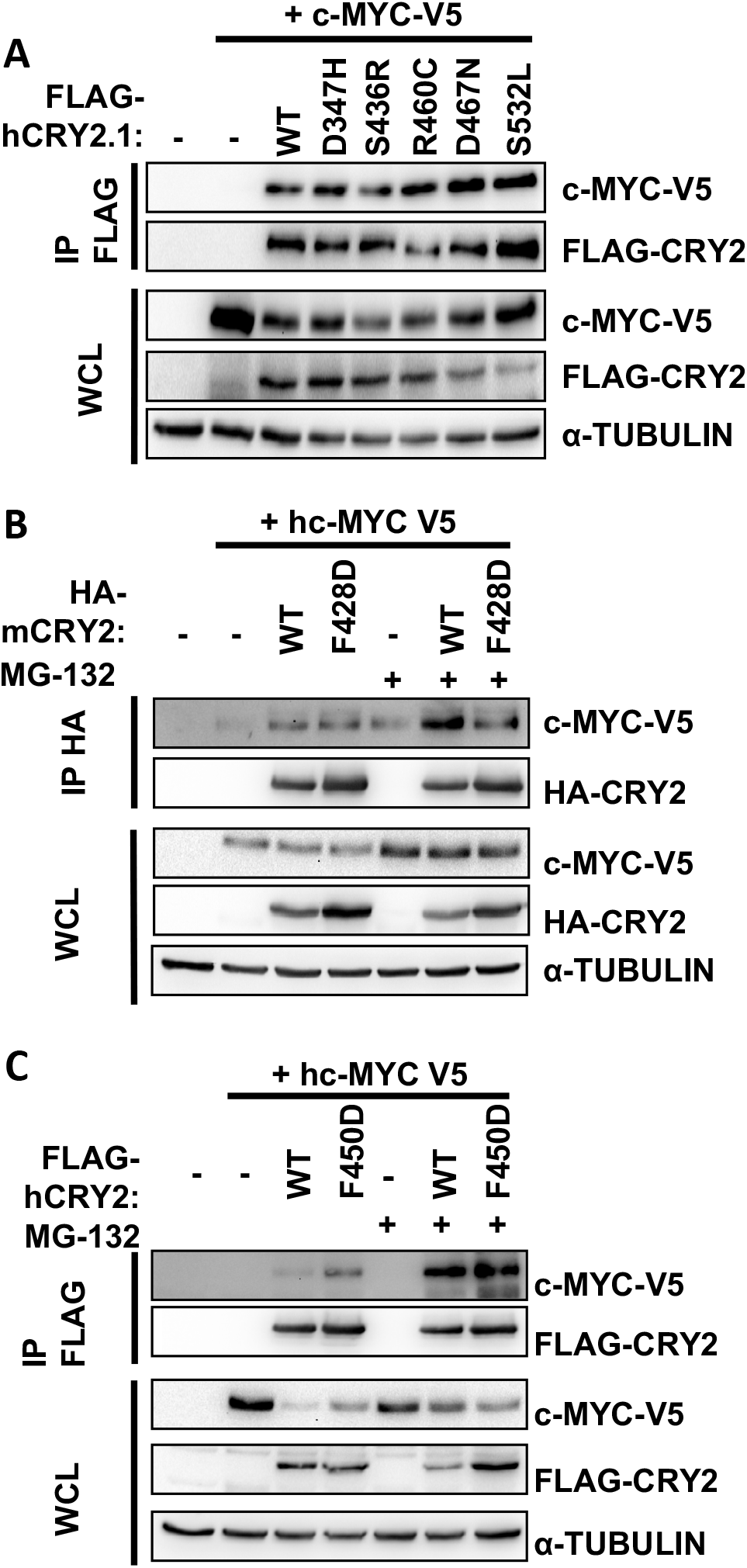
(A-C) Proteins detected by immunoblot (IB) following IP of FLAG (A,C) or HA (B) or in whole cell lysates (WCL) from HEK293T cells expressing the indicated plasmids with the indicated tags. For (B,C) samples were treated with vehicle (DMSO) or 10μM MG-132 (+) for four hours.

**Figure 2 - Figure Supplement 3.**
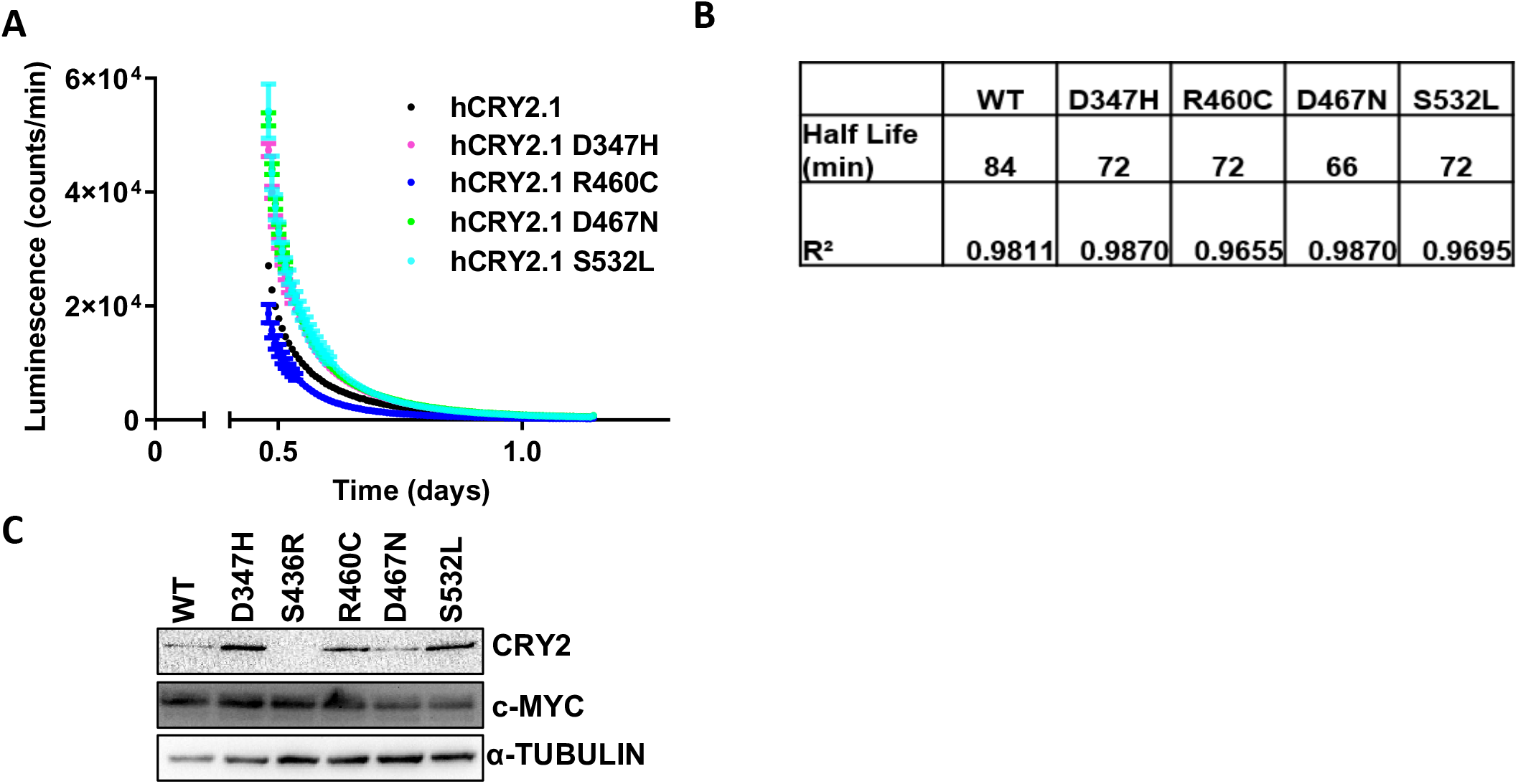
(A) Raw luminescence curves from one of three independent experiments in Figure 2C represented as mean ± SEM. (B) Half-lives (minutes) and R^2^ values are indicated from one-phase exponential curve fitting of data in (A) using Graph Pad Prism 9. (C) Western blots of lysates from cells in (A) blotted for the indicated proteins.

**Figure 3 - Figure Supplement 1.**
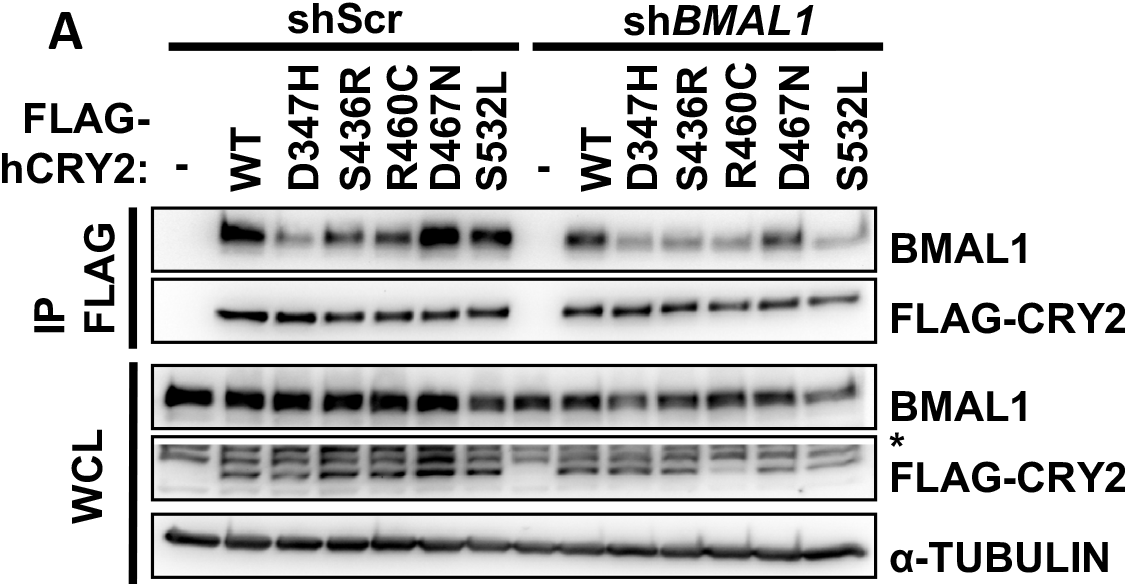
(A) Proteins detected by immunoblot (IB) following IP of the FLAG tag or in whole cell lysates (WCL) from HEK293T cells expressing the indicated plasmids with either knockdown using a control “scrambled” shRNA (shScr) or shRNA targeting Bmal1 (shBMAL1). * denotes non-specific band.

**Figure 3 - Figure Supplement 2.**
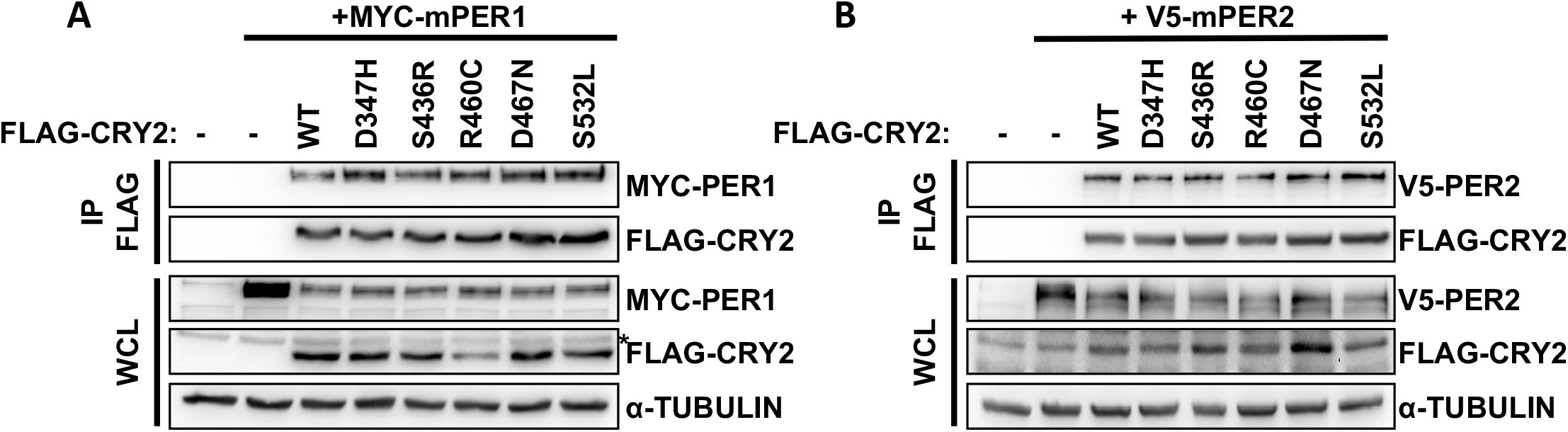
(A,B) Proteins detected by immunoblot (IB) following IP of the FLAG tag or in whole cell lysates (WCL) from HEK293T cells expressing the indicated plasmids. * denotes non-specific band.

**Figure 4 - Figure Supplement 1.**
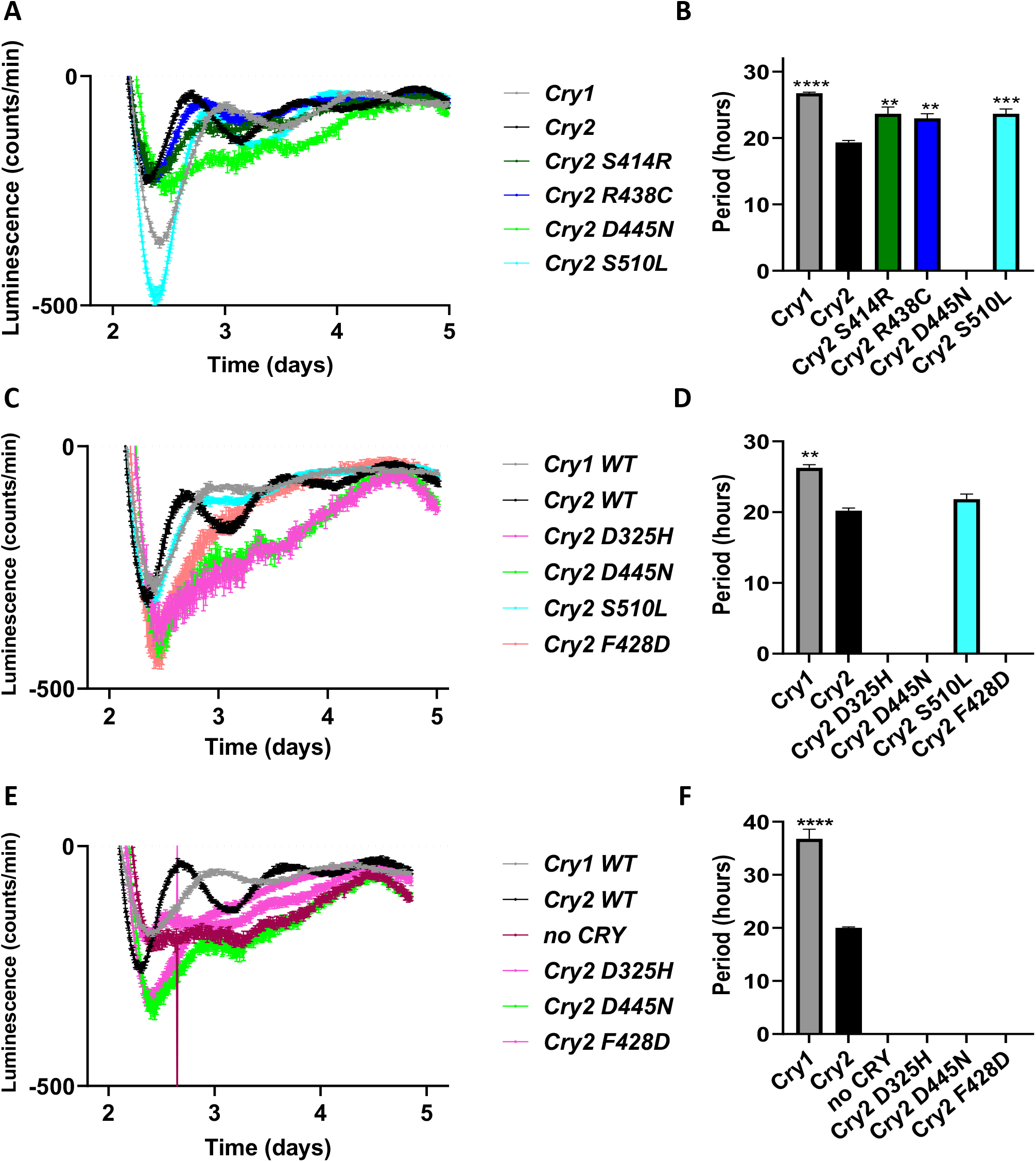
(A-F) Raw luminescence traces and periods from Figure 4D-F grouped as they were performed together. **p≤0.01, ***p≤0.001, ****p≤0.0001 by two-way ANOVA with Tukey’s multiple comparisons versus Cry2 in GraphPad Prism 9.

**Figure 5 - Figure Supplement 1.**
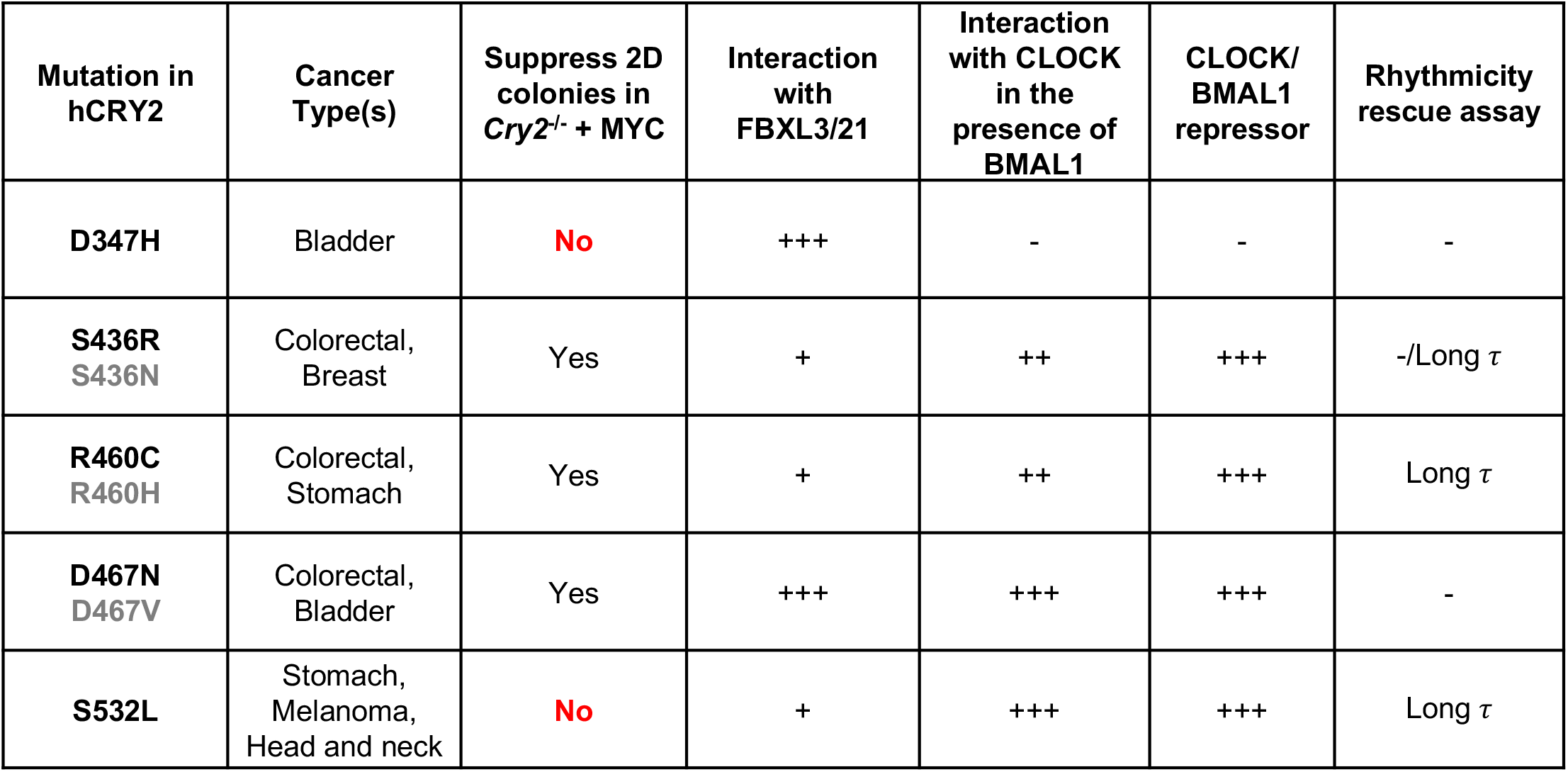
Table summarizing data from this paper. The first column shows point mutants that we studied in detail in black font and other mutations observed at the same location in grey font. For columns four and five, the number of +’s indicates strength of interaction and - indicates no detectable interaction as determined by co-IP. For column six, + indicates repressor of CLOCK/BMAL1 in a dose dependent manner in a luciferase assay, -indicates no repression of CLOCK/BMAL1. For column seven, -indicates arrhythmic; τ, period.

**Figure 5 - Figure Supplement 2.**
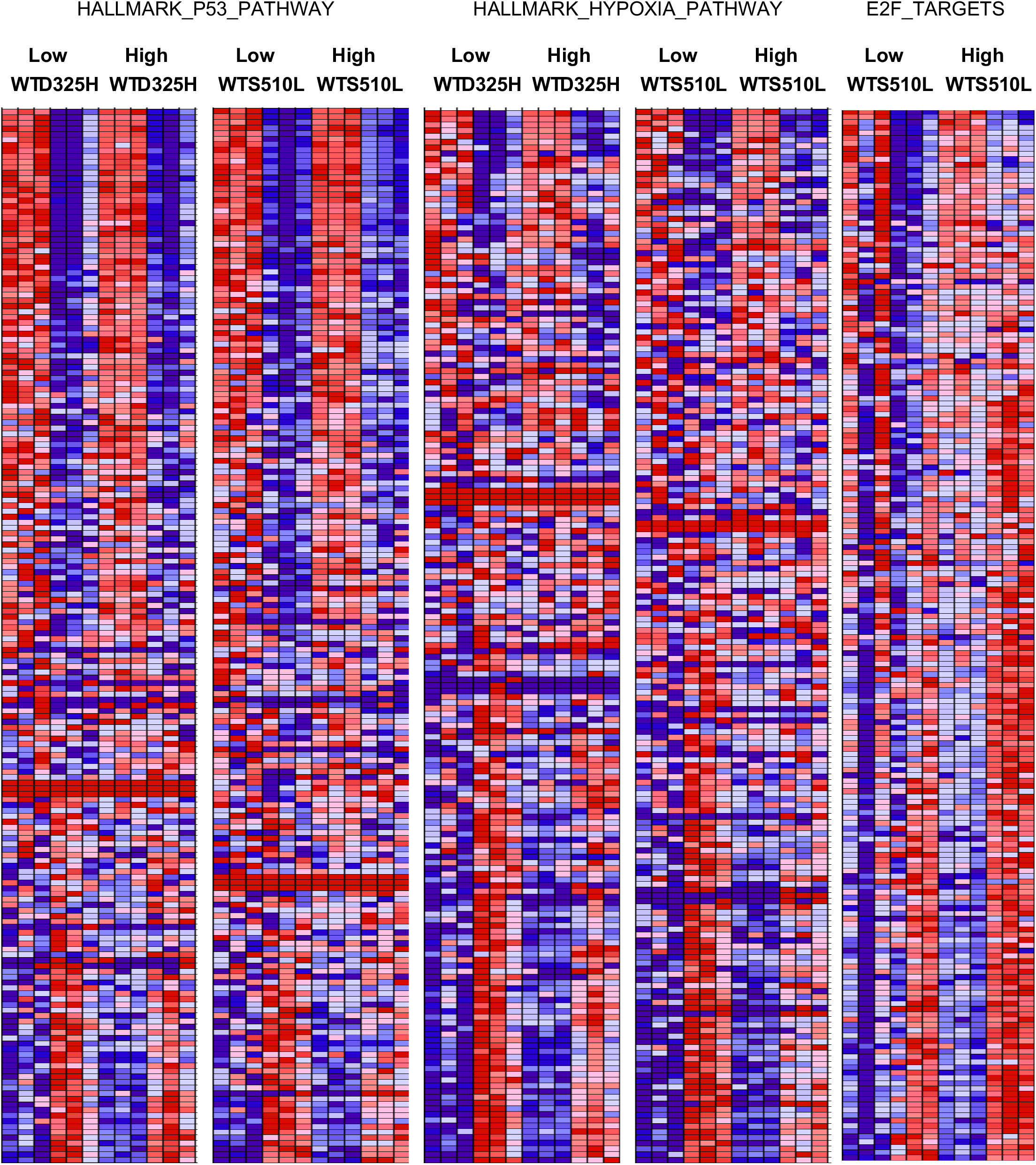
Heatmaps from RNA-sequencing representing all transcripts included in each gene set. Red, high expression; Blue low expression.

